# Evaluating the Robustness of Path-Preservation Benchmarks for Dimensionality Reduction Across Point-Density Thresholds: Linear and Cyclic Single-Cell Trajectories

**DOI:** 10.64898/2026.09.01.748633

**Authors:** Polina Bombina, Kevin R. Coombes

## Abstract

**Motivation:** Comparative studies of trajectory inference (TI) methods evaluate complete computational pipelines, making it impossible to isolate how much distortion is introduced specifically by the dimensionality reduction (DR) step. To our knowledge, no study has directly and systematically evaluated how well DR methods alone preserve a known reference path when projecting high-dimensional single-cell data to two dimensions, and no current study has introduced a dedicated set of metrics to qualify the degree of path-preservation quality after dimensionality reduction. This gap matters because DR is a universal preprocessing choice that shapes all downstream trajectory analysis, yet its independent geometric effect on path structure remains uncharacterised, and practitioners have no principled way to quantify it.

**Methods:** We tested a panel of candidate path-preservation metrics on two single-cell datasets with known reference trajectories — one linear, one cyclic — to determine whether the resulting metric values, DR-method rankings, and overall conclusions are sensitive to the number of points used to construct and display the path, and whether they remain stable once that choice is fixed. The primary, linear dataset is a CD4^+^ T-cell surface-protein dataset (3,096 cells, 51 proteins); a ground-truth reference path was constructed from cells lying close to the first principal component (PC1) of a single cluster, providing a known linear trajectory in the high-dimensional space. Sixteen DR methods were applied and twelve geometric path-preservation metrics were computed, spanning log-ratio distortions of length, curvature, and spatial similarity; Spearman rank correlations of pairwise distances and segment lengths; and structural complexity measures including self-intersection frequency and coiling. To test the sensitivity of this evaluation framework to path density, we varied the fraction of cells used to define the reference path from 1% to 10% (31–310 path points) and tracked how method rankings responded. The same analysis was repeated on a topologically distinct reference — a closed B-cell cell-cycle loop detected by persistent homology in a separate CyTOF dataset — to test whether these conclusions about metric and method stability hold for cyclic as well as linear trajectories.

**Results:** The central sensitivity question — whether the number of points used to construct the reference path changes the evaluation’s conclusions — was answered negatively on both datasets. On the linear PC1 trajectory, absolute values of all twelve metrics shifted smoothly as the path-density threshold was varied from 1% to 10% (31–310 points), reflecting the broadening of the reference band, but each method’s composite rank remained stable across every threshold: no method changed performance tier as the hyperparameter varied. A composite rank aggregating all twelve metrics identified the same consistently high-performing methods (UMAP, MDS, CNPE, TSNE, SPE, LPMIP) and consistently low-performing methods (SPMDS, LPP, DVE, LAPEIG, PHATE) at every density level tested. Considered on its own, `SpatDistSpear` — the single most discriminating metric — separated a high-fidelity group (LPMIP, DM, MDS, SPMDS, DVE, CISOMAP, CNPE; all *r* > 0.80) from a mid-range group (LAPEIG, SPE, PHATE, UMAP, TSNE, FOSMOD) and a low-fidelity group (PFA, NNP, LPP); global distance preservation and overall composite performance therefore do not always agree on the same “top tier” of methods, but this disagreement in *which* metric identifies the best methods is itself density-independent rather than an artefact of the specific threshold chosen. The cyclic loop reproduced the same density-independence: absolute metric values drifted with the per-segment band width, yet each method’s composite rank again held constant across all eleven density levels. The identity of the best and worst performers was largely, though not entirely, conserved between the two topologies — CNPE, LPMIP, SPE, and MDS were top performers and LAPEIG, DVE, and SPMDS were poor performers on both the linear path and the closed loop — with UMAP and TSNE the exception, dropping from top performers on the linear path to the middle of the sixteen-method panel, not the worst tier, on the closed loop. This topology-dependence is a property of the reference geometry rather than of path density: it holds consistently regardless of how many points are used to define the path.

**Significance:** This work introduces a direct, pipeline-independent evaluation of how DR methods distort trajectory geometry — a benchmarking dimension absent from existing TI comparisons. The within-dataset rank stability result, demonstrated on both a linear and a cyclic reference trajectory, validates the use of a fixed reference-path threshold as a robust operating point for large-scale DR benchmarking; however, the partial reordering of top performers between topologies shows that a method’s DR benchmark ranking is trajectory-shape-dependent and should not be assumed to transfer from a linear to a cyclic reference.

## 1 Introduction

Single-cell proteomics and transcriptomics generate high-dimensional feature matrices in which biologically meaningful transitions — differentiation, activation, maturation — manifest as continuous trajectories through cell space. A central challenge in exploratory data analysis is whether these trajectories can be faithfully recovered after dimensionality reduction (DR), which collapses the high-dimensional representation to two or three dimensions for visualisation and downstream inference.

Numerous DR methods have been proposed, differing in their mathematical objectives: some optimise global distance preservation (MDS, CISOMAP), others prioritise local neighbourhood structure (UMAP, t-SNE, LAPEIG), and others model diffusion processes or graph-based connectivity (Diffusion Maps, PHATE, LPP). The degree to which each method distorts a known reference trajectory is therefore expected to vary, and quantitative benchmarking is needed to guide method selection.

Our broader framework evaluates path preservation using twelve geometric metrics applied to a reference path constructed from cells lying close to the first principal component of the data. A key free parameter in this construction is the fraction of cells included: a tight threshold selects only the most linearly arranged cells, while a loose threshold introduces off-axis cells that may add curvature. We evaluate this hyperparameter choice on two topologically distinct reference trajectories. The primary analysis uses a linear reference path derived from the PC1 axis of a CD4^+^ T-cell cluster. To assess whether the findings generalise beyond linear structure, we repeat the sensitivity analysis on a closed cyclic trajectory: a B-cell cell-cycle loop identified by persistent homology in a separate CyTOF dataset. Together, these two references — one linear, one cyclic — allow us to determine whether the hyperparameter choice affects the relative ranking of DR methods, or only the absolute scale of the metrics, and to test whether that conclusion is robust across different trajectory topologies.

## 2 Methods

### 2.1 Reference Trajectory Design

We analyzed two complementary reference trajectory structures: a linear trajectory derived from the CAVA CD4 T-cell dataset and a cyclic trajectory derived from a B-cell CyTOF cell-cycle dataset.

#### 2.1.1 Linear Trajectory Construction

For the linear trajectory analysis, we selected a biologically coherent subpopulation from the CAVA dataset. Specifically, **cluster 5** (3096 cells × 51 proteins), identified in the UMAP projection from Seurat, was used for all downstream analyses. Protein expression values were **mean-centered but not scaled** prior to analysis. This preprocessing preserves the original Euclidean distance relationships among observations, which is critical for assessing geometric distortions introduced by DR methods. In contrast, scaling would alter relative distances and thus confound interpretation of preservation metrics.

Principal component analysis (PCA) was performed in the original 51-dimensional protein space. The **first principal component (PC1)**, representing the dominant axis of variation, was used to define a linear trajectory. To construct a reference path in the high-dimensional space, we quantified each cell’s alignment with PC1 by computing its **orthogonal (perpendicular) distance** to the PC1 axis. Cells with small perpendicular distances lie close to this axis and therefore approximate a one-dimensional trajectory.

Cells were selected using **quantile-based thresholding**. For a given threshold *t*, we retained the fraction of cells with the smallest perpendicular distances to PC1, producing an ordered subset that approximates a continuous path. This approach generalizes prior fixed-threshold definitions by enabling systematic variation in path density.

To assess robustness of DR methods with respect to path definition, we evaluated ten density levels ranging from 1% to 10% of cells closest to PC1. Lower thresholds yield sparse trajectories tightly constrained to the principal axis, whereas higher thresholds incorporate increasingly peripheral cells, introducing greater thickness, curvature, and potential geometric complexity.

#### 2.1.2 Cyclic Trajectory Construction

To complement the linear trajectory with a topologically distinct structure, we analyzed a B-cell CyTOF dataset in which cell-cycle progression forms a closed loop. The loop was identified using persistent homology via a Vietoris–Rips filtration, and the most persistent one-dimensional cycle was selected as the reference trajectory.

The resulting loop consists of 15 nodes (cells) connected by 15 edges, forming a closed path without a defined start or end point. Accordingly, all downstream evaluation metrics were adapted to account for the cyclic topology. The same set of 16 DR methods was applied to this dataset to enable direct comparison with the linear trajectory analysis.

To vary path density, we employed a **per-segment densification strategy**. For each of the 15 loop edges, we retained the *q*% of cells closest to that segment and then took the union across all segments. The density parameter was evaluated over the same range as in the linear case, from 1% to 10%, allowing direct comparison of DR method robustness across linear and cyclic trajectory topologies.

### 2.2 DR Methods

Sixteen dimensionality reduction methods were evaluated. UMAP was computed using the umap R package, Diffusion Maps using the diffusionMap package, and all remaining methods were implemented using the Rdimtools package.

Each constructed path was then mapped into lower-dimensional projections.

### 2.3 Path Preservation Metrics

Twelve independent metrics were tracked to map structural deformations across the threshold continuum. These mathematical criteria assess different aspects of geometric warping and are categorized by their optimization targets:

1. **Log-Ratio Deviations (Target** → **0):** Tracks metric scale variations where an optimal score of zero shows perfect geometric preservation symmetry (`LenDistort`, `CurvRatio`, `SegVarRatio`, `SpatSimRatio`, `EndpointDisp`).
2. **Rank Optimization Metrics (Target** → **1):** Non-parametric Spearman correlation measures evaluating monotonic trajectory agreement and alignment accuracy (`CurvSpear`, `SegLenSpear`, `SpatDistSpear`).
3. **Structural Complexity Metrics:** Geometric descriptors of the embedded path’s shape. `Smoothness`, `Contact`, and `CrossingNorm` penalize localized high-frequency coordinate variance, micro-oscillations, and self-intersections (ideal = low / 0), whereas `Coil` measures the spatial extent of the path and is reported with ideal = high.

All metrics are computed by the Preservation package.

## 3 Results

### 3.1 CAVA CD4 T-cell cluster (linear trajectory)

Cluster 5 of the CAVA CD4 T-cell dataset comprises 3,096 cells profiled across 51 surface proteins. Figure 1A shows the global UMAP embedding of all sequenced cells coloured by Seurat cluster identity, confirming that Cluster 5 forms a spatially coherent region in the embedding. Figure 1B shows the PCA projection of Cluster 5 alone, with cells partitioned into four immunophenotypically distinct sub-populations by Mercator hierarchical clustering. The elongated arrangement of sub-populations along PC1, rather than discrete separation, supports the interpretation of PC1 as a continuous reference trajectory through the cluster.

**Figure 1:**
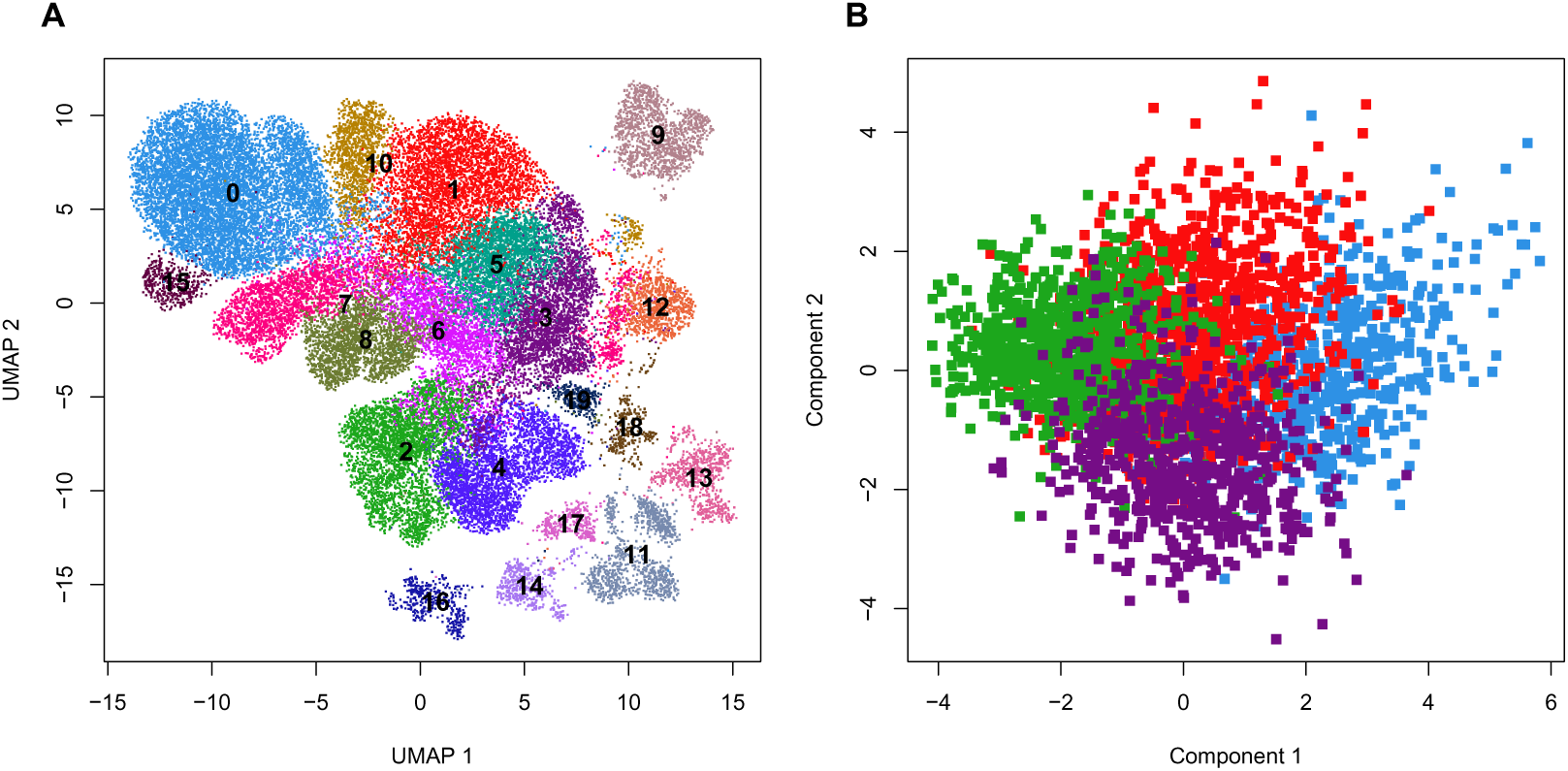
Topological layout of CD4+ T-cell clusters. (A) Global UMAP visualization of all sequenced cells colored by Seurat cluster identities. (B) Principal Component Analysis (PCA) projection of target Cluster 5, partitioned into four immunophenotypically distinct sub-populations by the Mercator R framework.

To construct a high-dimensional reference path, we computed for every cell its perpendicular distance to the PC1 axis — that is, the Euclidean norm of all PC scores excluding PC1. Figure 2 shows the resulting distribution of projection residuals across the 3,096 cells. The distribution is right-skewed: a small fraction of cells lie very close to PC1 while the majority are distributed at progressively greater radial distances. Vertical lines mark the 5th through 20th percentiles of this distribution. This structure confirms that a meaningful linear substructure exists in the protein space and that tightening the quantile threshold selects an increasingly pure, geometrically coherent approximation of the PC1 trajectory.

**Figure 2:**
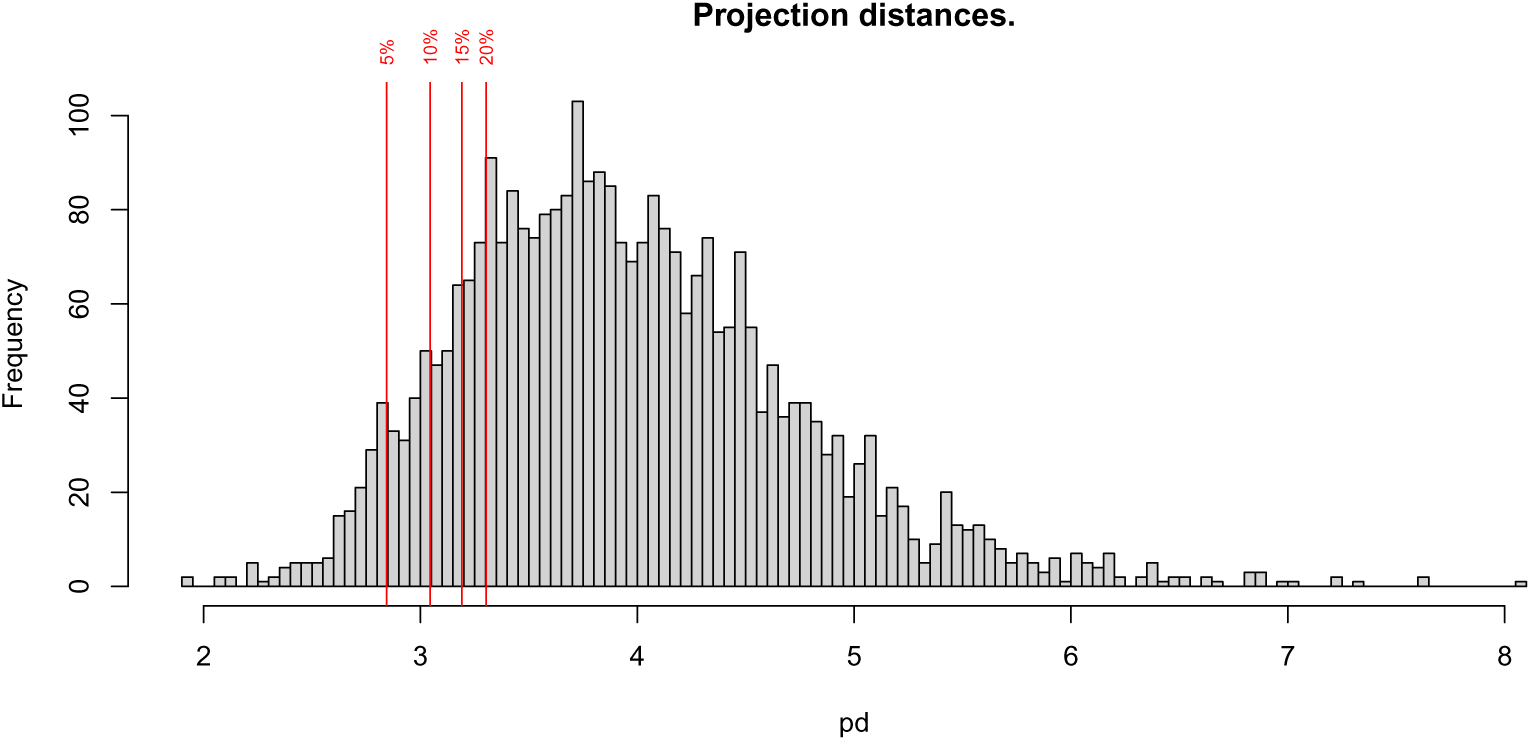
Distribution of perpendicular distances from PC1, with 5–20% quantile thresholds.

For each threshold *t* ∈ {1%, 2%*, . . .,* 10%}, the ⌊*t* × 3096⌋ cells with the smallest perpendicular residuals were retained and ordered by their PC1 score to define the reference path. At the 10% threshold this yields 310 cells, spanning the full extent of the principal trajectory. Lower thresholds produce sparser but more strictly linear paths, while higher thresholds introduce cells farther from PC1, adding curvature and broadening the path band.

Sixteen DR methods were applied to the full cluster. Figure 3 displays the resulting 16 two-dimensional projections, coloured by Mercator sub-cluster identity. Substantial qualitative differences in layout are immediately apparent: spectral methods (LAPEIG, DVE, SPMDS) spread sub-clusters into extended, ribbon-like arrangements; neighbour-based methods (UMAP, TSNE) form compact, separated clusters; and manifold-interpolation methods (LPP, SPE) produce irregular, diffuse configurations.

**Figure 3:**
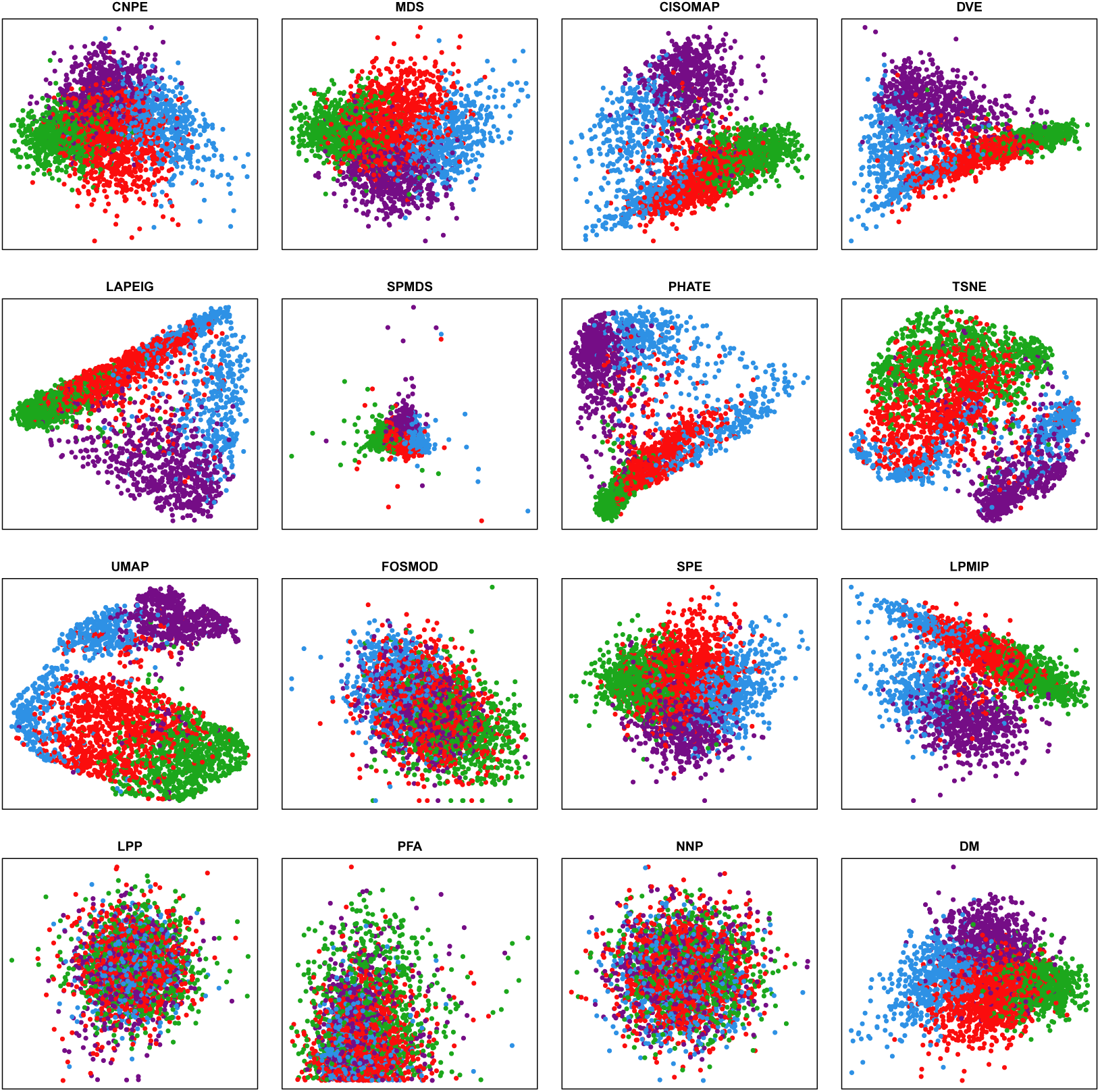
2D projections of all cluster cells, one panel per DR method.

Figure 4 overlays the reference path on each embedding at the 10% threshold. Grey points show the full set of cluster cells; the black line traces the ordered path; the green dot marks the PC1-minimum endpoint and the red dot the PC1-maximum endpoint. Visual inspection reveals pronounced method-level differences in how the ordered path traverses the embedding space. Methods such as DVE and LAPEIG map the path into an elongated ribbon that spans the full extent of the embedding, while UMAP and TSNE compress it into a densely overlapping tangle within a compact cluster region. LPP and SPE produce highly disordered paths with frequent back-crossings. SPMDS collapses the entire path to a near-point. The per-metric sensitivity curves (Figure 5) allow visual tracking of how these patterns change as the cell band widens from 31 cells (1%) to 310 cells (10%). In all cases the qualitative method ordering is preserved; methods that disperse the path widely at 1% continue to do so at 10%.

**Figure 4:**
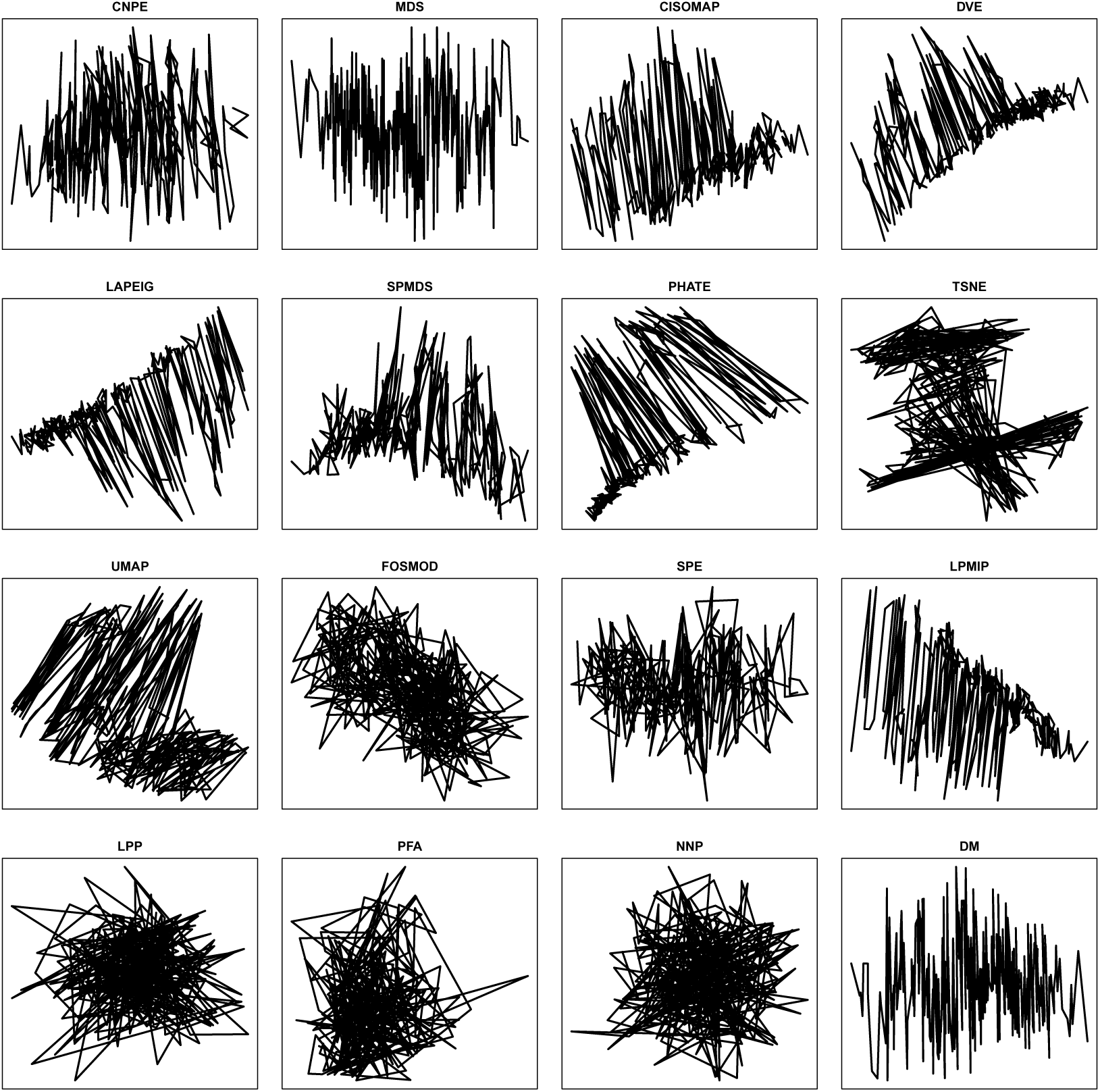
The reference path projected into each projection (10% threshold).

**Figure 5:**
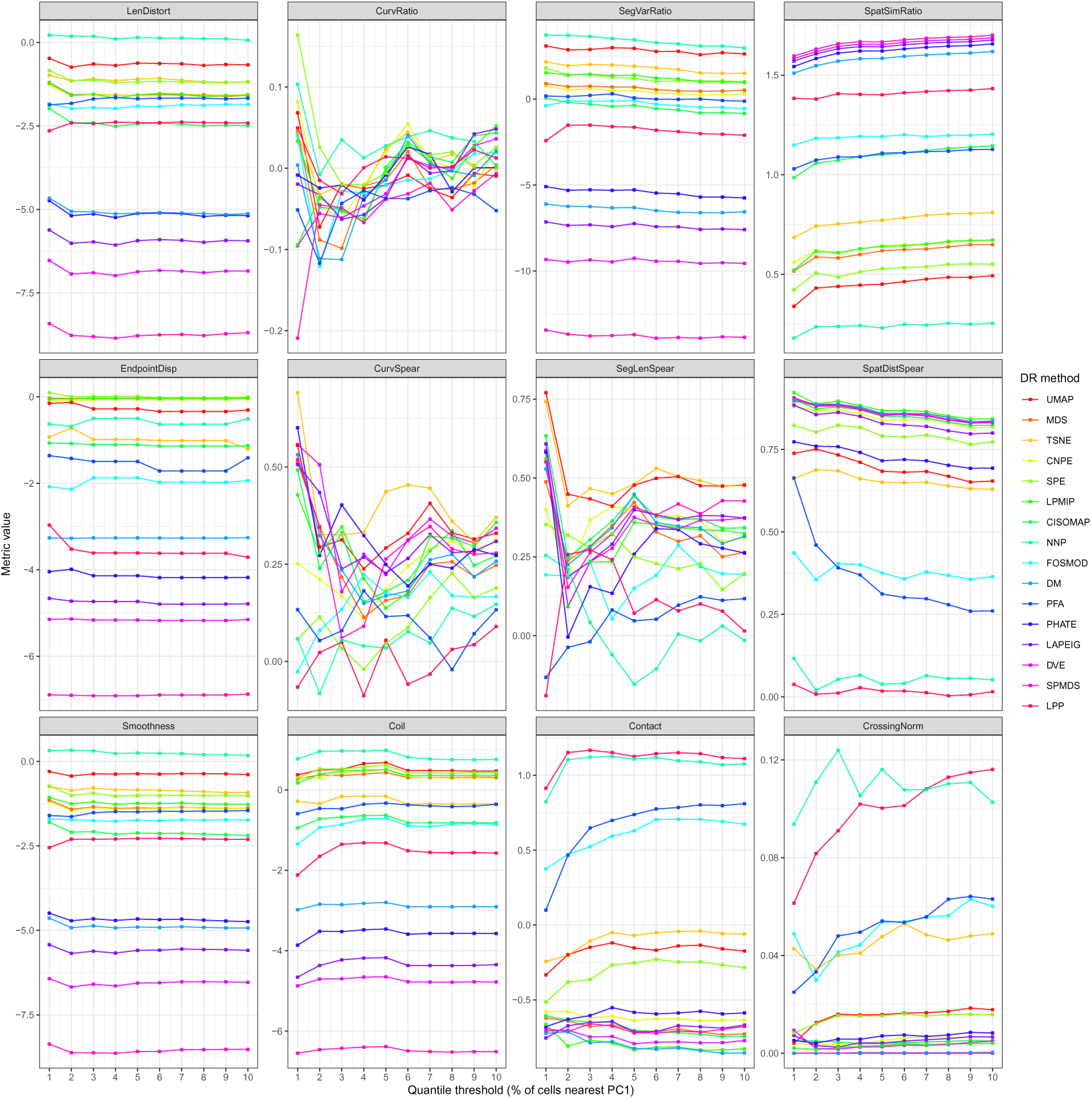
Per-metric sensitivity curves. Each panel shows one metric; each line is one DR method. The x-axis shows the quantile threshold as a percentage of cells. Most metrics show parallel shifts with stable relative ordering; CrossingNorm is nearly invariant.

#### 3.1.1 Quantitative path preservation at the 10% reference threshold

Twelve preservation metrics were computed for all 16 DR methods using the 310-cell reference path (10% threshold). Table 2 presents the full metric matrix; key patterns are summarised below.

**Table 1:** Dimension reduction algorithms used in the study (chrono-logical order).

| Abbreviation | Meaning | Year | Study |
| --- | --- | --- | --- |
| MDS | MultiDimensional Scaling | 1964 | Kruskal (1964) |
| CISOMAP | Conformal Isometric Feature Mapping | 2003 | Silva & Tenenbaum (2003) |
| LAPEIG | Laplacian Eigenmaps | 2003 | Belkin & Niyogi (2003) |
| LPP | Locality Preserving Projection | 2003 | He & Niyogi (2003) |
| SPE | Stochastic Proximity Embedding | 2004 | Agrafiotis (2003) |
| DM | Diffusion Maps | 2005 | Nadler et al. (2005) |
| PFA | Principal Feature Analysis | 2007 | Lu et al. (2007) |
| TSNE | t-distributed Stochastic Neighbor Embedding | 2008 | van der Maaten & Hinton (2008) |
| DVE | Distinguishing Variance Embedding | 2009 | Qinggang et al. (2009) |
| CNPE | Complete Neighborhood Preserving Embedding | 2010 | Wang et al. (2010) |
| NNP | Nearest Neighbor Projection | 2010 | Tejada et al. (2003) |
| LPMIP | Locality-Preserved Maximum Information Projection | 2011 | Wang et al. (2011) |
| SPMDS | Spectral Multidimensional Scaling | 2013 | Aflalo & Kimmel (2013) |
| FOSMOD | Forward Orthogonal Search by Maximizing the Overall Dependency | 2013 | Senawi et al. (2017) |
| UMAP | Uniform Manifold Approximation and Projection | 2018 | McInnes et al. (2018) |
| PHATE | Potential of Heat Diffusion for Affinity-based Transition Embedding | 2019 | Moon et al. (2019) |

**Table 2:**
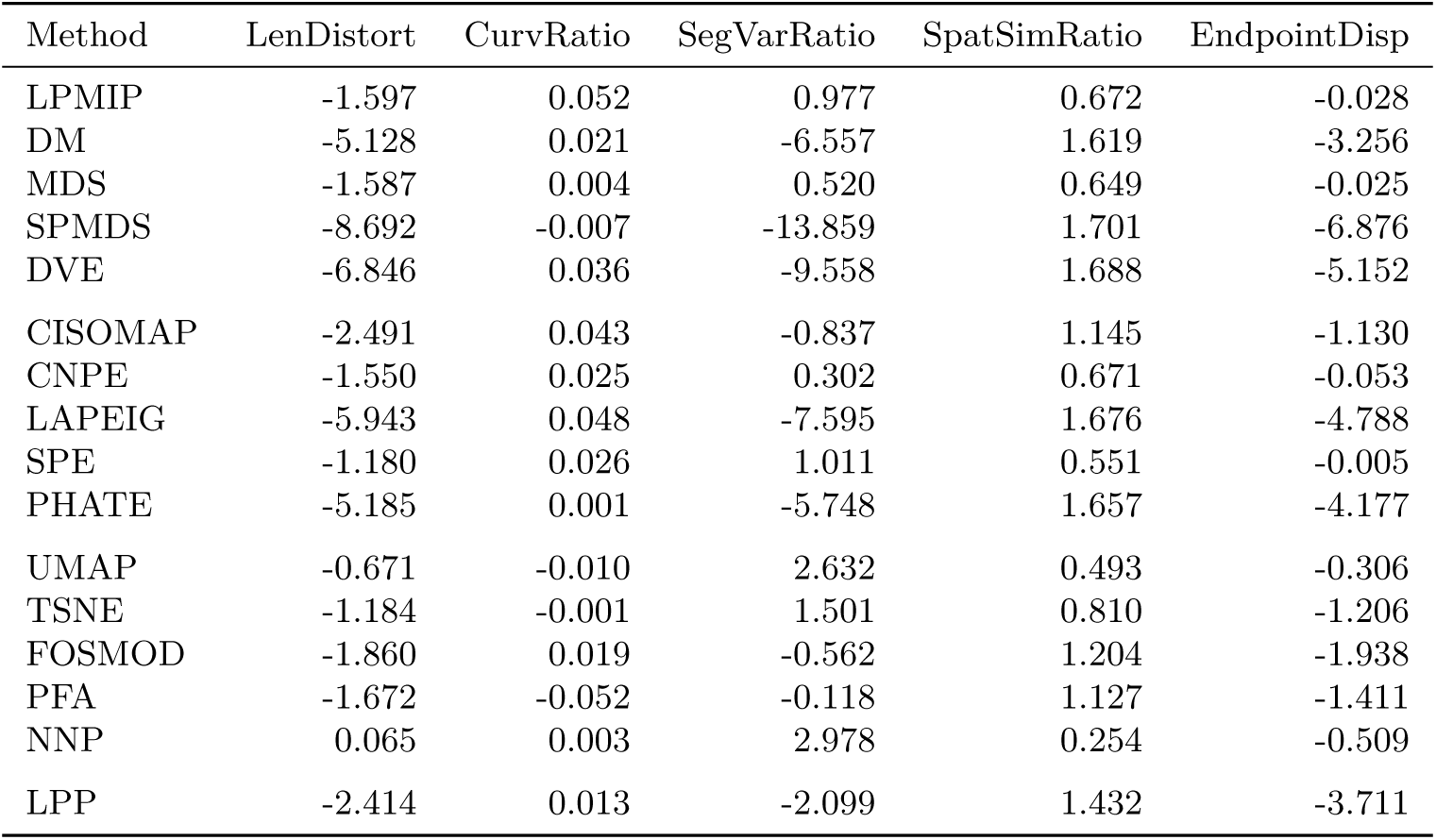
Path-preservation metrics at the 10% threshold (n = 310). Part A: log-ratio metrics (ideal = 0).

| Method | LenDistort | CurvRatio | SegVarRatio | SpatSimRatio | EndpointDisp |
| --- | --- | --- | --- | --- | --- |
| LPMIP | -1.597 | 0.052 | 0.977 | 0.672 | -0.028 |
| DM | -5.128 | 0.021 | -6.557 | 1.619 | -3.256 |
| MDS | -1.587 | 0.004 | 0.520 | 0.649 | -0.025 |
| SPMDS | -8.692 | -0.007 | -13.859 | 1.701 | -6.876 |
| DVE | -6.846 | 0.036 | -9.558 | 1.688 | -5.152 |
| CISOMAP | -2.491 | 0.043 | -0.837 | 1.145 | -1.130 |
| CNPE | -1.550 | 0.025 | 0.302 | 0.671 | -0.053 |
| LAPEIG | -5.943 | 0.048 | -7.595 | 1.676 | -4.788 |
| SPE | -1.180 | 0.026 | 1.011 | 0.551 | -0.005 |
| PHATE | -5.185 | 0.001 | -5.748 | 1.657 | -4.177 |
| UMAP | -0.671 | -0.010 | 2.632 | 0.493 | -0.306 |
| TSNE | -1.184 | -0.001 | 1.501 | 0.810 | -1.206 |
| FOSMOD | -1.860 | 0.019 | -0.562 | 1.204 | -1.938 |
| PFA | -1.672 | -0.052 | -0.118 | 1.127 | -1.411 |
| NNP | 0.065 | 0.003 | 2.978 | 0.254 | -0.509 |
| LPP | -2.414 | 0.013 | -2.099 | 1.432 | -3.711 |

**Log-ratio metrics** (ideal value = 0) revealed pronounced compression across all methods: `LenDistort` was negative for 15 of 16 methods, indicating that path length is systematically reduced in the lower-dimensional space, with SPMDS (−8.69) and LAPEIG (−5.94) showing the most extreme compression and NNP (+0.07) and UMAP (−0.67) the least. `CurvRatio` was close to zero for all methods (range −0.05 to +0.05), indicating that mean local turning angle is approximately preserved regardless of method. `SegVarRatio` varied widely: CNPE (+0.30) and MDS (+0.52) remained close to zero while SPMDS (−13.82) and LAPEIG (−7.60) showed extreme variance reduction. `SpatSimRatio` was positive for all methods, reflecting an increase in mean pairwise spatial similarity in the 2D embedding.

**Rank-correlation metrics** (ideal value = 1) discriminated methods most clearly on SpatDistSpear, the Spearman correlation of pairwise distances between the high- and low-dimensional paths. Three distinct performance tiers emerged: a high-fidelity group of seven methods comprising LPMIP (0.843), DM (0.836), MDS (0.835), SPMDS (0.834), DVE (0.831), CISOMAP (0.824), and CNPE (0.818), all strictly exceeding *r* = 0.80; a mid-range group including LAPEIG (0.800), SPE (0.773), PHATE (0.693), UMAP (0.654), TSNE (0.629), and FOSMOD (0.365); and a low-fidelity group comprising PFA (0.260), NNP (0.051), and LPP (0.016), for which global pairwise distance relationships are effectively destroyed in the 2D embedding.

**Structural complexity metrics** reinforced these tiers. `CrossingNorm` was zero for MDS, SPMDS, and DM — their projected paths contain no self-intersections — while LPP (0.116) and NNP (0.103) exhibited the highest rates of self-crossing. `Smoothness` and `Coil`, as assessed by the `Preservation` package functions, showed SPMDS, DVE, and DM having the largest-magnitude values, consistent with their extended, ribbon-like spatial distributions.

#### 3.1.2 Sensitivity of preservation metrics to path density

Figure 5 shows the value of each of the 12 metrics as a function of the percentage threshold, sweeping from 1% (31 cells) to 10% (310 cells). To facilitate cross-threshold comparison, Table 4 summarises the mean and maximum absolute change between the 1% and 10% thresholds for each metric, and Tables 5–6 give the per-method standard deviation of each metric across the ten threshold levels.

**Table 3:** Path-preservation metrics at the 10% threshold (n = 310). Part B: Spearman metrics (ideal = 1); Smoothness and Contact (ideal = low); Coil (ideal = high); CrossingNorm (ideal = 0).

| Method | CurvSpear | SegLenSpear | SpatDistSpear | Smoothness | Coil | Contact | CrossingNorm |
| --- | --- | --- | --- | --- | --- | --- | --- |
| LPMIP | 0.357 | 0.324 | 0.843 | -1.274 | 0.359 | -0.828 | 0.004 |
| DM | 0.258 | 0.316 | 0.836 | -4.928 | -2.908 | -0.855 | 0.000 |
| MDS | 0.247 | 0.265 | 0.835 | -1.385 | 0.314 | -0.729 | 0.000 |
| SPMDS | 0.279 | 0.427 | 0.834 | -8.522 | -6.508 | -0.678 | 0.000 |
| DVE | 0.343 | 0.374 | 0.831 | -6.537 | -4.775 | -0.772 | 0.005 |
| CISOMAP | 0.272 | 0.342 | 0.824 | -2.193 | -0.820 | -0.743 | 0.005 |
| CNPE | 0.313 | 0.309 | 0.818 | -1.388 | 0.291 | -0.634 | 0.007 |
| LAPEIG | 0.309 | 0.373 | 0.800 | -5.591 | -4.346 | -0.668 | 0.007 |
| SPE | 0.189 | 0.197 | 0.773 | -1.011 | 0.424 | -0.284 | 0.016 |
| PHATE | 0.273 | 0.262 | 0.693 | -4.742 | -3.572 | -0.587 | 0.008 |
| UMAP | 0.330 | 0.477 | 0.654 | -0.392 | 0.470 | -0.174 | 0.018 |
| TSNE | 0.371 | 0.480 | 0.629 | -0.917 | -0.352 | -0.060 | 0.049 |
| FOSMOD | 0.167 | 0.195 | 0.365 | -1.731 | -0.856 | 0.673 | 0.060 |
| PFA | 0.133 | 0.117 | 0.260 | -1.454 | -0.359 | 0.810 | 0.063 |
| NNP | 0.147 | -0.015 | 0.051 | 0.174 | 0.758 | 1.075 | 0.103 |
| LPP | 0.090 | 0.015 | 0.016 | -2.308 | -1.570 | 1.111 | 0.116 |

**Table 4:**
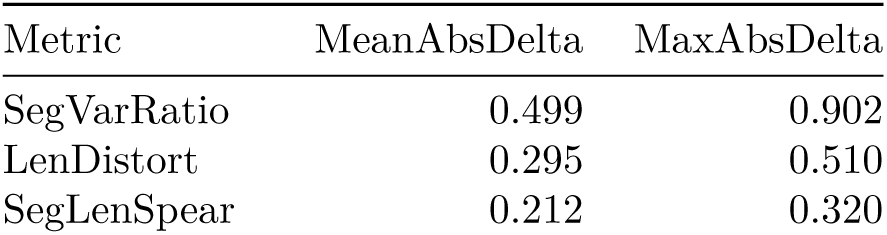
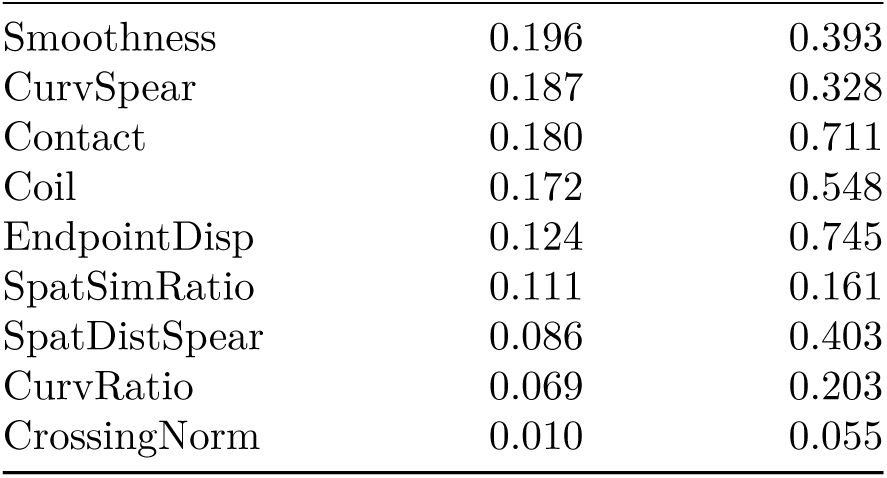
Mean and maximum absolute change per metric between the 1% and 10% thresholds, across all 16 methods.

**Table 5:** Per-method SD across the 1%–10% grid — Part A: log-ratio metrics. Methods sorted by overall sensitivity (most stable first).

| Method | LenDistort | CurvRatio | SegVarRatio | SpatSimRatio | EndpointDisp |
| --- | --- | --- | --- | --- | --- |
| CNPE | 0.095 | 0.044 | 0.174 | 0.034 | 0.013 |
| FOSMOD | 0.052 | 0.044 | 0.181 | 0.016 | 0.088 |
| UMAP | 0.069 | 0.035 | 0.163 | 0.045 | 0.078 |
| MDS | 0.120 | 0.044 | 0.151 | 0.040 | 0.004 |
| NNP | 0.046 | 0.030 | 0.277 | 0.022 | 0.074 |
| DVE | 0.121 | 0.043 | 0.098 | 0.032 | 0.013 |
| LAPEIG | 0.120 | 0.038 | 0.147 | 0.032 | 0.045 |
| TSNE | 0.067 | 0.027 | 0.238 | 0.038 | 0.118 |
| DM | 0.144 | 0.052 | 0.183 | 0.033 | 0.004 |
| SPMDS | 0.120 | 0.057 | 0.144 | 0.032 | 0.012 |
| LPMIP | 0.124 | 0.046 | 0.208 | 0.045 | 0.008 |
| SPE | 0.107 | 0.053 | 0.251 | 0.040 | 0.035 |
| PFA | 0.074 | 0.027 | 0.139 | 0.030 | 0.144 |
| PHATE | 0.143 | 0.021 | 0.221 | 0.034 | 0.066 |
| LPP | 0.077 | 0.022 | 0.295 | 0.017 | 0.210 |
| CISOMAP | 0.155 | 0.041 | 0.296 | 0.048 | 0.026 |

**Table 6:** Per-method SD across the 1%–10% grid — Part B: Spearman and shape metrics.

| Method | CurvSpear | SegLenSpear | SpatDistSpear | Smoothness | Coil | Contact | CrossingNorm |
| --- | --- | --- | --- | --- | --- | --- | --- |
| CNPE | 0.067 | 0.067 | 0.022 | 0.070 | 0.083 | 0.023 | 0.001 |
| FOSMOD | 0.074 | 0.060 | 0.026 | 0.019 | 0.177 | 0.116 | 0.010 |
| UMAP | 0.087 | 0.100 | 0.036 | 0.033 | 0.091 | 0.061 | 0.004 |
| MDS | 0.112 | 0.079 | 0.022 | 0.078 | 0.048 | 0.037 | 0.000 |
| NNP | 0.065 | 0.128 | 0.025 | 0.055 | 0.106 | 0.090 | 0.008 |
| DVE | 0.116 | 0.125 | 0.026 | 0.070 | 0.069 | 0.033 | 0.002 |
| LAPEIG | 0.089 | 0.113 | 0.029 | 0.071 | 0.137 | 0.033 | 0.002 |
| TSNE | 0.114 | 0.090 | 0.020 | 0.056 | 0.090 | 0.072 | 0.005 |
| DM | 0.109 | 0.086 | 0.023 | 0.089 | 0.053 | 0.051 | 0.000 |
| SPMDS | 0.130 | 0.107 | 0.024 | 0.079 | 0.056 | 0.021 | 0.000 |
| LPMIP | 0.089 | 0.119 | 0.025 | 0.062 | 0.094 | 0.052 | 0.001 |
| SPE | 0.079 | 0.064 | 0.020 | 0.090 | 0.091 | 0.090 | 0.002 |
| PFA | 0.056 | 0.083 | 0.124 | 0.060 | 0.077 | 0.224 | 0.013 |
| PHATE | 0.117 | 0.154 | 0.029 | 0.070 | 0.111 | 0.034 | 0.001 |
| LPP | 0.061 | 0.136 | 0.010 | 0.082 | 0.231 | 0.074 | 0.017 |
| CISOMAP | 0.097 | 0.103 | 0.026 | 0.111 | 0.100 | 0.049 | 0.000 |

The sensitivity curves (Figure 5, Table 4) show that SegVarRatio carries the largest absolute shift between 1% and 10% (mean |Δ| = 0.50, max 0.90), followed by LenDistort (mean 0.29). Crucially, however, these shifts are largely parallel across methods: the rank ordering within each metric panel is preserved even when the absolute values drift. `CrossingNorm` is essentially invariant (mean |Δ| = 0.01), and `CurvRatio` also shows minimal change (mean 0.07). The three Spearman metrics decline gently with increasing threshold, reflecting that wider bands include cells farther from the linear PC1 axis — a property of the reference path rather than the embeddings.

Figure 6 shows the overall sensitivity of each method to the threshold choice, defined as the mean rescaled standard deviation across all 12 metrics. CNPE is the most threshold-stable method (sensitivity score 0.27), followed by FOSMOD (0.30) and UMAP (0.34). The most threshold-sensitive methods are CISOMAP (0.52), LPP (0.51), and PHATE (0.45). Notably, threshold-sensitivity does not align simply with quality: LPP is both a poor performer and highly sensitive, while CISOMAP is a strong overall performer that is nonetheless among the most threshold-sensitive. The sensitivity reflects in part which aspects of the path geometry change most as the band widens.

**Figure 6:**
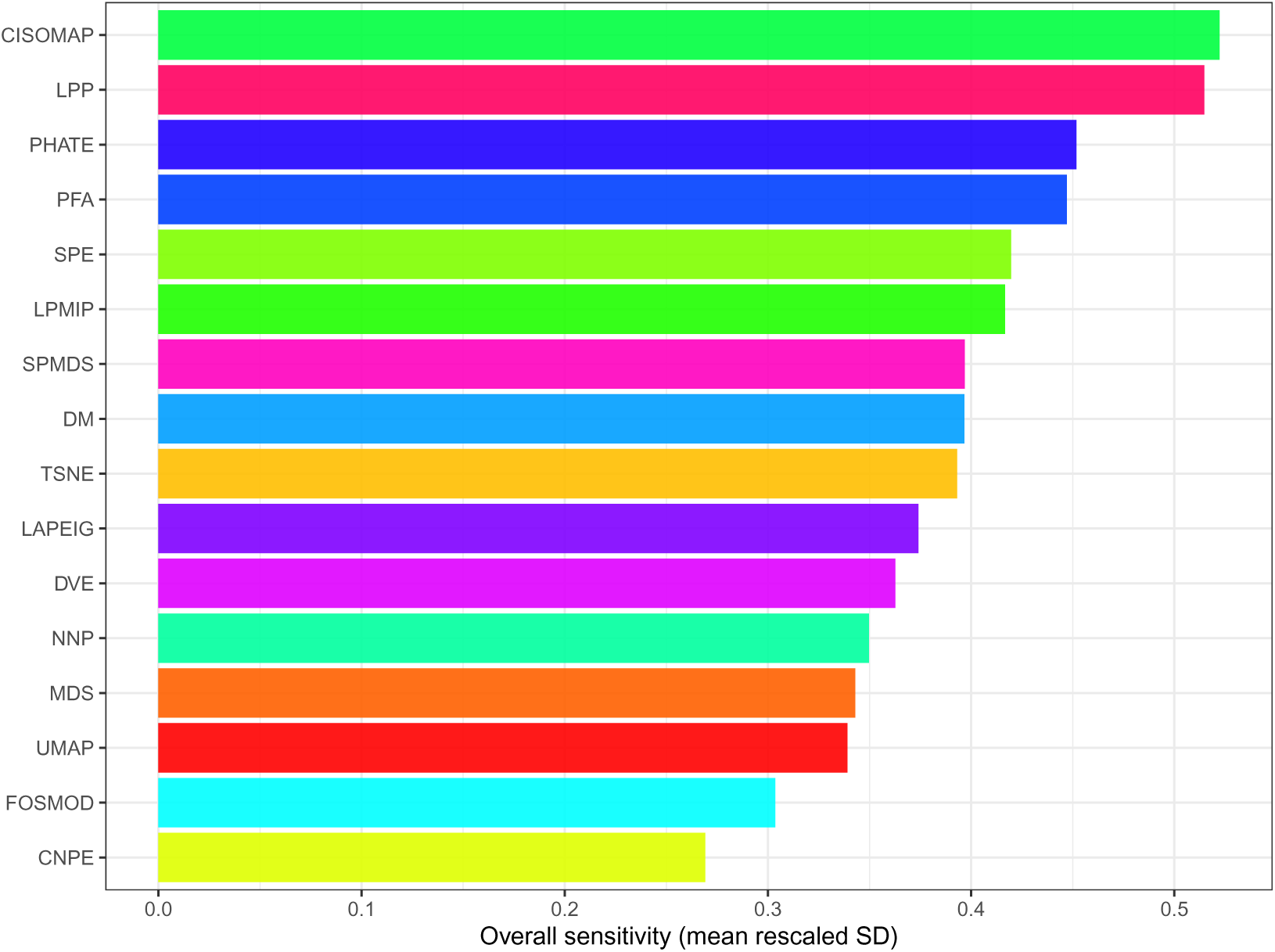
Overall sensitivity of each method to the percentage threshold, measured as mean rescaled standard deviation across the 12 metrics. Lower values indicate more threshold-stable behaviour. Bar colours match each method’s colour in Figures 5 and 7.

#### 3.1.3 Non-parametric rank stability across thresholds

To assess directly whether the performance conclusions about DR methods depend on the threshold, we assigned each method a direction-adjusted dense rank on each of the 12 metrics at each threshold level, then computed a composite rank as the mean rank across all available metrics. Dense ranks (ties assigned to the same integer without gaps) were used to produce clean, comparable scores.

The density-rank pattern across all metrics and thresholds is summarised in the beanplot below (Figure 7). Methods with narrow, sharply concentrated beans are threshold-stable; methods with wide beans vary more across metrics and thresholds. UMAP, MDS, TSNE, and CNPE are concentrated at low rank values (good performance, stable). SPMDS, LPP, DVE, and LAPEIG are concentrated at high rank values (poor performance, also stable). The width of each bean reflects cross-metric variation — no method is uniformly best or worst across all twelve metrics — but the overall composite ordering is clear and persistent. Notably, UMAP and TSNE appear near the top of the beanplot despite mid-range SpatDistSpear values (0.654 and 0.629 respectively); this reflects their favourable scores on structural complexity metrics such as CrossingNorm and Contact, which pull their composite rank upward relative to their global-distance performance alone.

**Figure 7:**
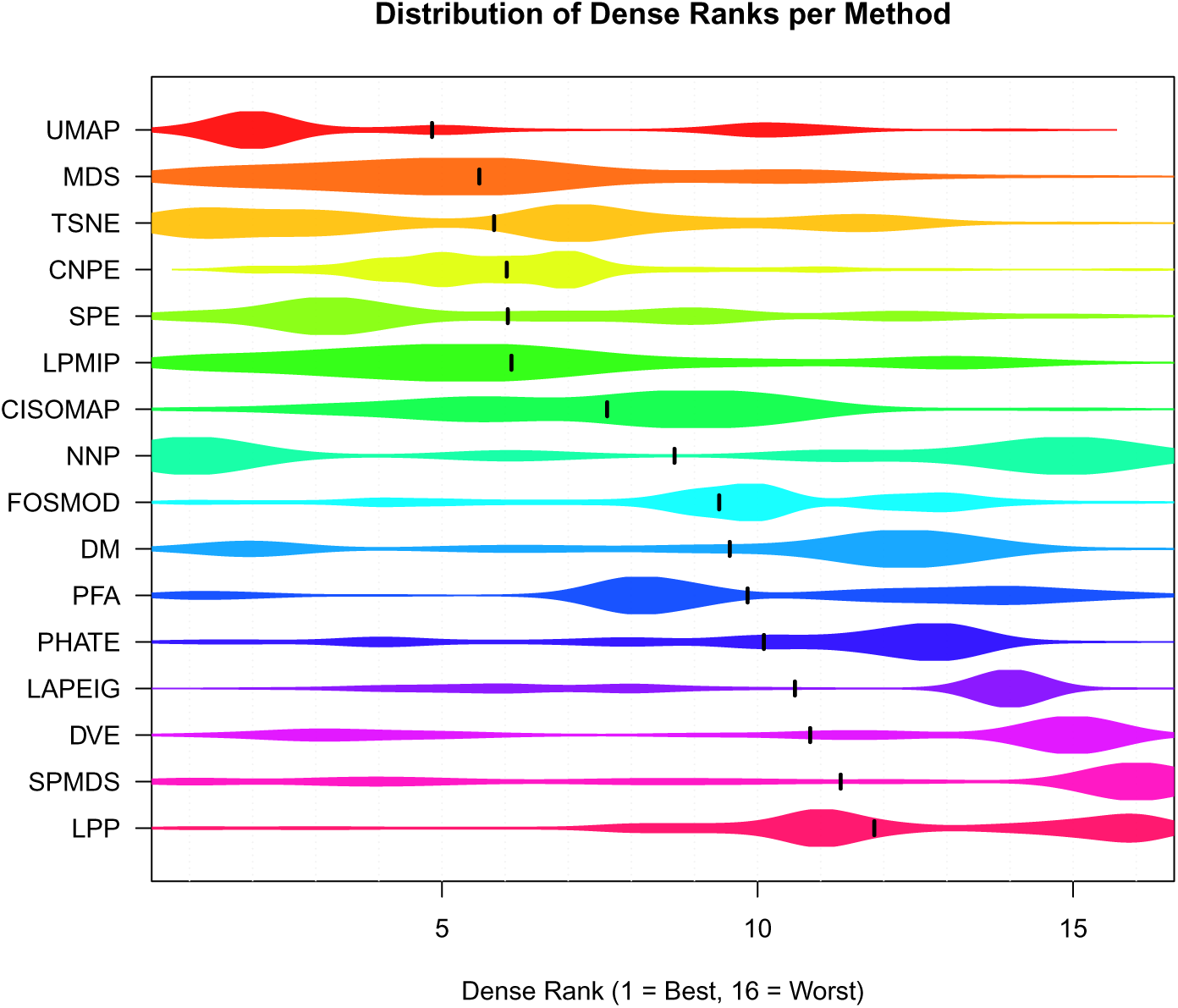
Distribution of composite dense ranks per method across all thresholds and all 12 metrics. Rank 1 is best, rank 16 is worst. Methods are sorted by their overall mean rank. The vertical bar in each bean marks the mean.

**Figure 8:**
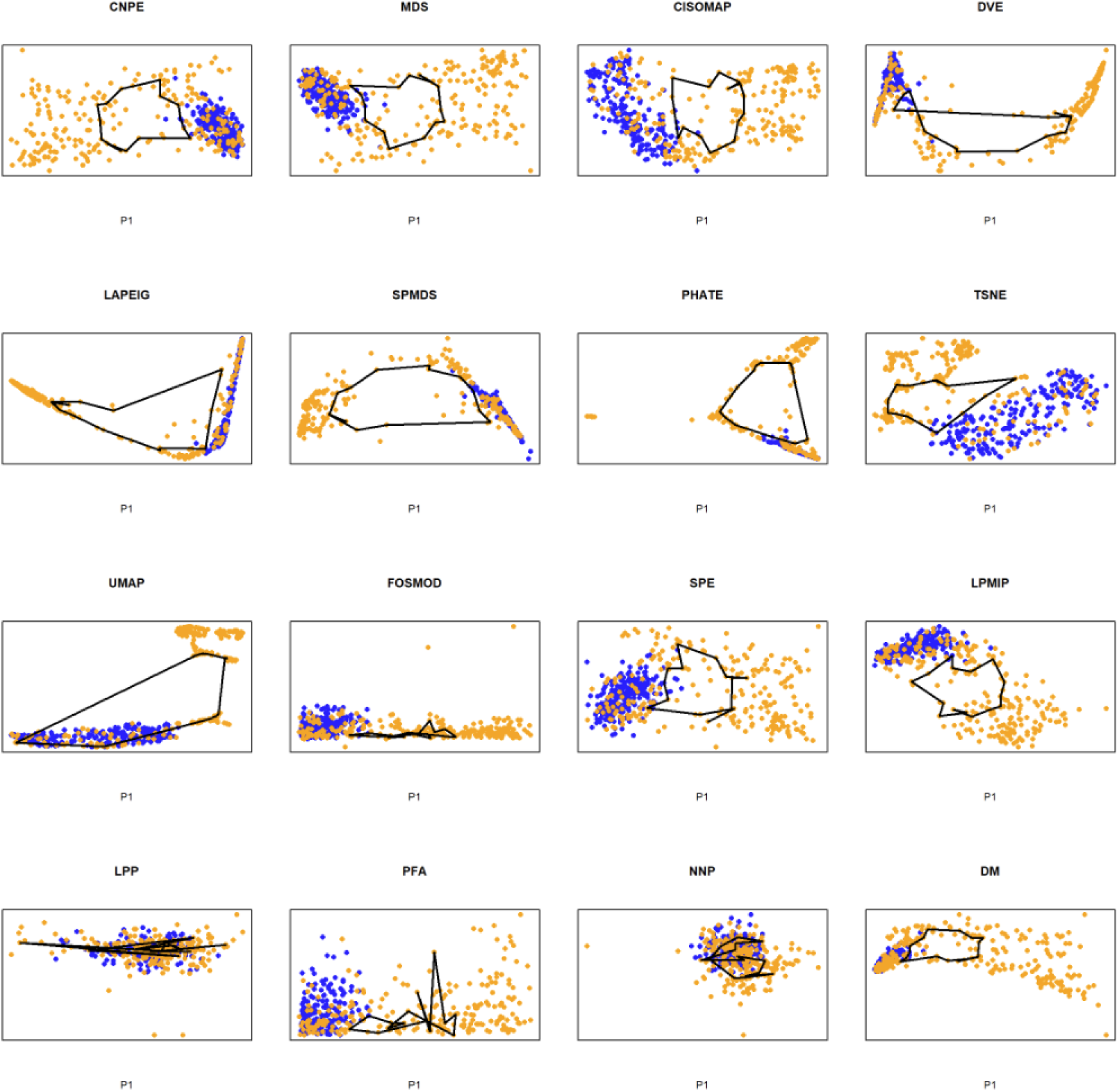
Cell cycle TDA loop.

**Figure 9:**
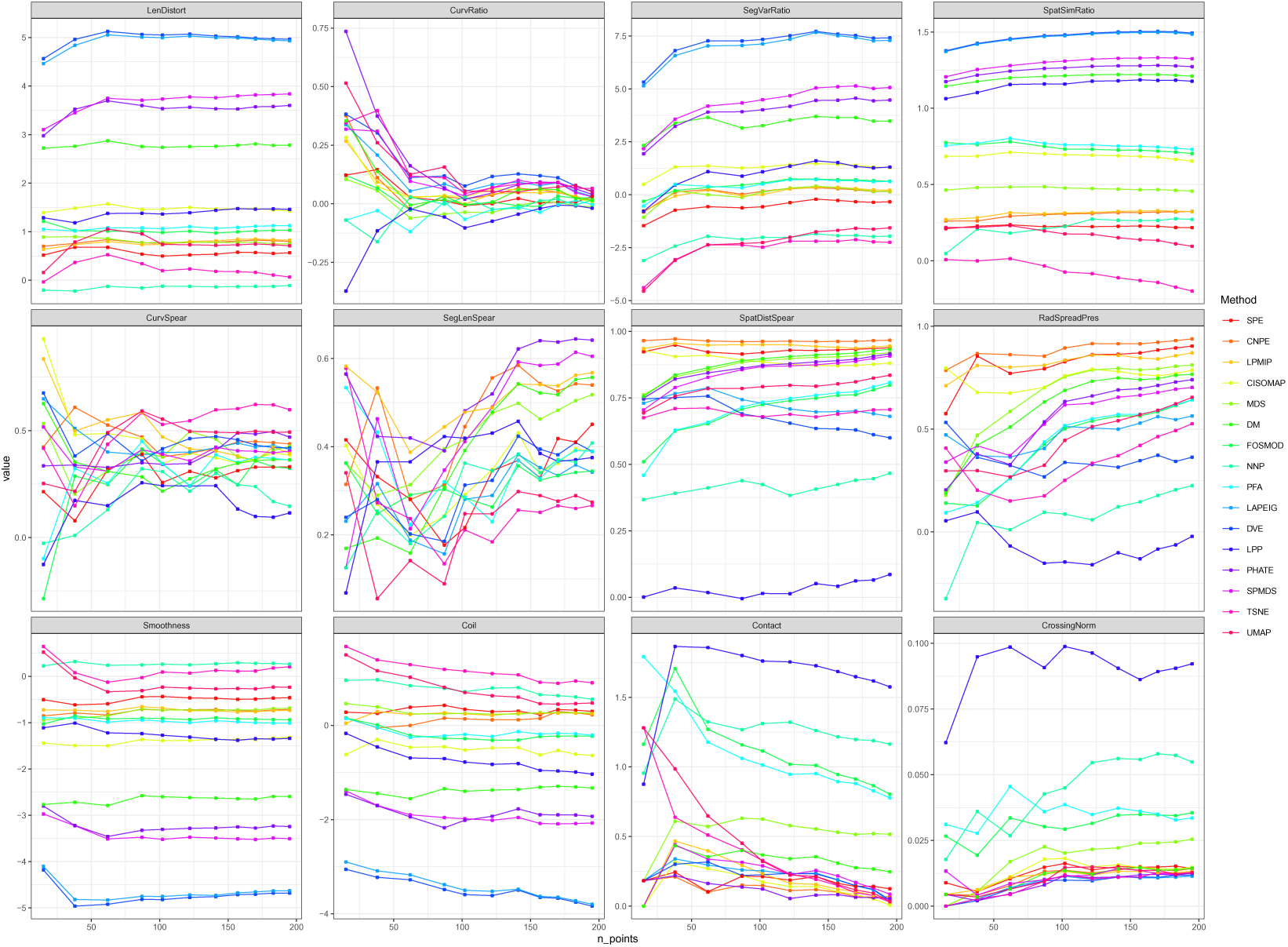
Each metric vs number of loop points, one line per DR method. The x-axis is the number of ordered loop points (cf. Figure 5, where the linear analysis uses the percentage threshold directly).

**Figure 10:**
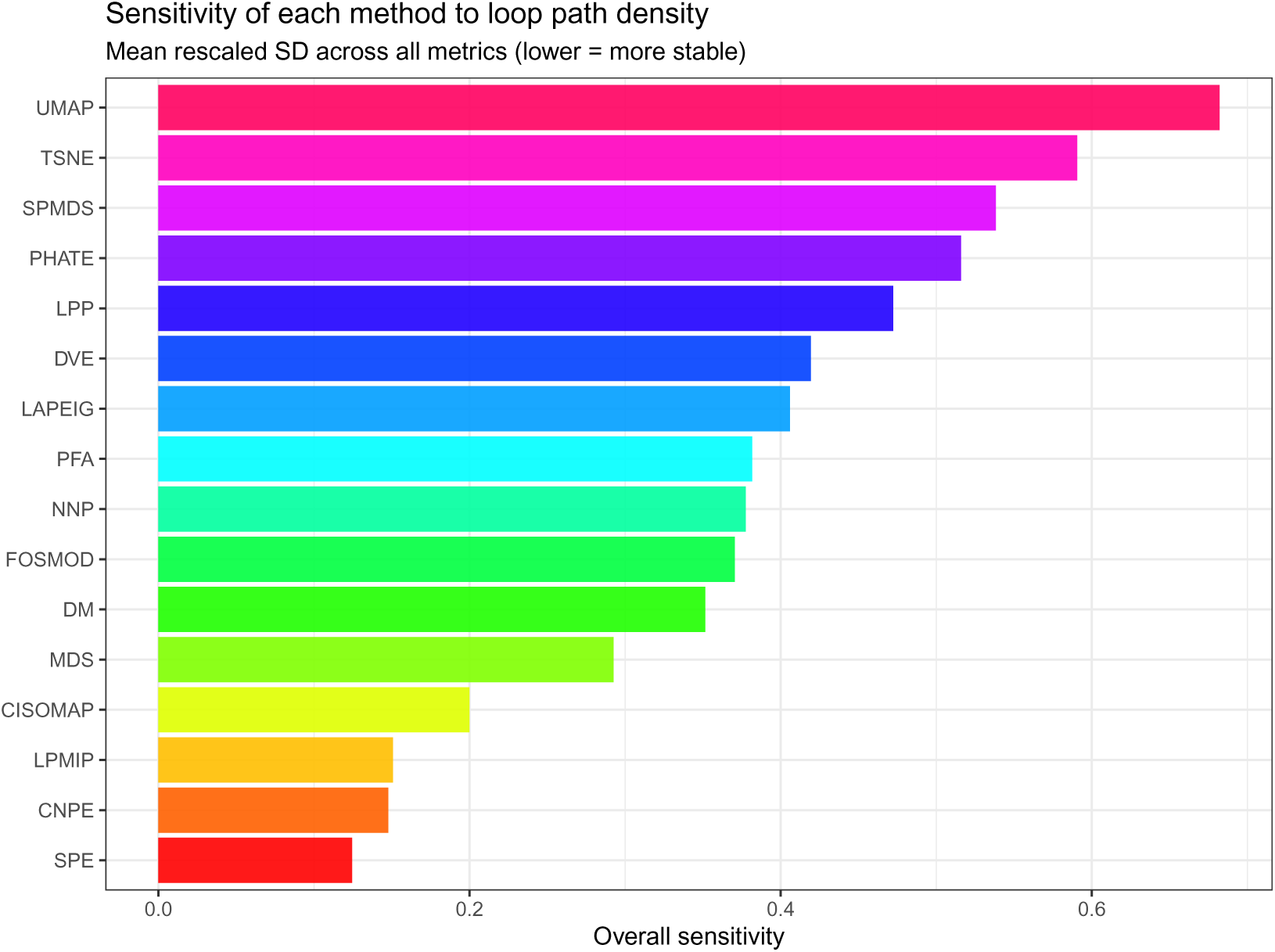
Per-method sensitivity to path density: mean rescaled SD across all metrics.

**Figure 11:**
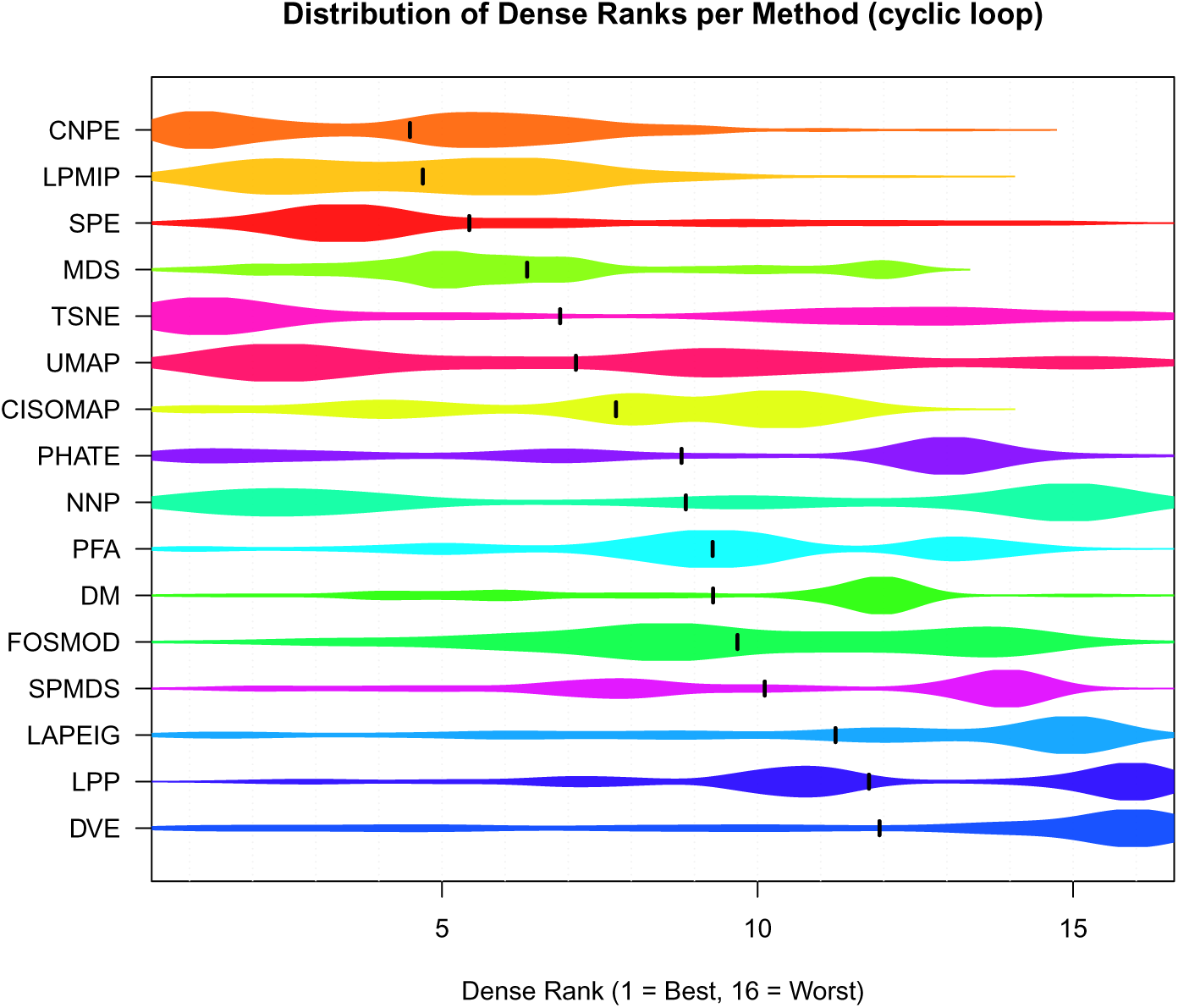
Distribution of composite dense ranks per method across all density levels and all 12 cyclic metrics. Rank 1 is best. Methods are sorted by overall mean rank; the vertical bar in each bean marks the mean. Colours match the cyclic curves and bar chart.

Taken together, Figures 6 and 7 demonstrate that the high-performing methods (UMAP, MDS, CNPE, TSNE, SPE, LPMIP) and the low-performing methods (SPMDS, LPP, DVE, LAPEIG, PHATE) retain their relative positions at every threshold examined. No method crosses from the top tier to the bottom tier, or vice versa, as the threshold is varied.

### 3.2 B-cell CyTOF dataset (cyclic trajectory)

The analysis above used a *linear* reference path (the PC1 trajectory of a CD4^+^ T-cell cluster). A natural question is whether the same robustness holds for a *closed* trajectory, where the reference is a loop rather than a line. To test this, we repeated the path-density sensitivity analysis on a B-cell CyTOF dataset whose cell-cycle structure forms a topological cycle, detected by persistent homology (TDA). This complements the linear case with a topologically distinct reference and a partly different metric set adapted to closed paths.

Because the reference is a closed loop, the metric set differs slightly from the linear case: `EndpointDisp` is undefined for a closed path and is dropped, leaving four log-ratio metrics (`LenDistort`, `CurvRatio`, `SegVarRatio`, `SpatSimRatio`), four Spearman preservation metrics — three of which (`CurvSpear`, `SegLenSpear`, `SpatDistSpear`) are computed by exactly the same `Preservation` package functions as in the linear analysis, plus the loop-specific `RadSpreadPres` — and four shape/structural metrics from the `Preservation` package (`Smoothness`, `Coil`, `Contact`, `CrossingNorm`) — twelve metrics in total, the same count as the linear analysis. All twelve, including the closed-loop log-ratio metrics, are computed by functions exported from the `Preservation` package rather than defined inline. The densification strategy also differs: the loop’s 15 nodes define 15 line segments, and for each segment we keep the *q*% of cells closest to it, unioned and ordered around the cycle. This gives even coverage around the loop. We retain the 15-node loop as baseline and sweep *q* from 1% to 10% in 1% steps, exactly as in the linear (CAVA) analysis.

~~~
## Performing eigendecomposition
## Computing Diffusion Coordinates
## Used default value: 6 dimensions
## Elapsed time: 0.12 seconds
~~~

The ordered TDA loop has 15 nodes and therefore 15 line segments (edges) joining consecutive nodes around the closed cycle. For each segment we compute every cell’s perpendicular distance to that segment, and keep the *q*% of cells closest to it. Taking the union across all 15 segments and ordering by arc-position around the loop gives the densified path. Because each segment selects its own closest cells, coverage is even all the way around the cycle — sparse regions of the loop are not starved relative to dense ones, which is the failure mode of a single global distance threshold. The 15-node loop is retained as the baseline, and *q* is swept from 1% to 10% as in the linear (CAVA) analysis.

The log-ratio metrics use the closed-loop (wraparound) variants CyclicLengthDistortion, CyclicCurvature, and `CyclicSegmentVariance`, since a loop has no start/end point and `diff()`-based segments would miss the edge that closes the cycle back on itself. `SpatSimRatio` uses the same `SpatialSimilarity` as the linear analysis, since it depends only on the set of pairwise distances and is unaffected by wraparound. The three Spearman preservation metrics (`CurvSpear`, `SegLenSpear`, `SpatDistSpear`) and the three shape metrics (`Smoothness`, `Coil`, `Contact`) are computed by exactly the same package functions used in the linear (CAVA) analysis. `RadSpreadPres` is the one loop-specific metric with no linear analogue — the Spearman correlation of each point’s distance from the path centroid, via `RadialSpreadPreservation`, which is meaningful only for a closed loop. `CrossingNorm`, the normalised self-intersection count of the projected loop, is computed by the same `Preservation::CrossingNorm` function used in the linear analysis, applied to the closed 2D path.

#### 3.2.1 Quantitative path preservation at the 15-node baseline

As for the linear path, we first report the full metric matrix for all methods, then examine sensitivity across density levels. Here the matrix is reported at the **15-node baseline loop** — the original persistent-homology cycle, before any densification — so the table reflects how each method preserves the bare topological loop. The metrics are split into two tables for readability, mirroring the linear-path presentation.

#### 3.2.2 Sensitivity of preservation metrics to path density

Unlike the linear sensitivity curves (Figure 5), where the x-axis is the percentage threshold, the cyclic curves below use the *number of ordered loop points* on the x-axis. This is because the per-segment densification produces a different number of retained cells at each level (Table 8), and plotting against the actual point count gives a more faithful picture of how the metrics respond to path density.

**Table 7:** Overall sensitivity of each DR method to the path-density threshold, ranked from most stable (rank 1) to least stable. Score = mean rescaled SD across all 12 metrics.

| Rank | Method | OverallSensitivity |
| --- | --- | --- |
| 1 | CNPE | 0.269 |
| 2 | FOSMOD | 0.304 |
| 3 | UMAP | 0.339 |
| 4 | MDS | 0.343 |
| 5 | NNP | 0.350 |
| 6 | DVE | 0.363 |
| 7 | LAPEIG | 0.374 |
| 8 | TSNE | 0.393 |
| 9 | DM | 0.397 |
| 10 | SPMDS | 0.397 |
| 11 | LPMIP | 0.417 |
| 12 | SPE | 0.420 |
| 13 | PFA | 0.447 |
| 14 | PHATE | 0.452 |
| 15 | LPP | 0.515 |
| 16 | CISOMAP | 0.522 |

**Table 8:** Density level and the resulting number of ordered loop points.

| Level | n_points |
| --- | --- |
| 15 nodes | 15 |
| 1% | 38 |
| 2% | 62 |
| 3% | 87 |
| 4% | 102 |
| 5% | 122 |
| 6% | 141 |
| 7% | 157 |
| 8% | 170 |
| 9% | 183 |
| 10% | 195 |

**Table 9:** Cyclic-loop preservation metrics at the 15 nodes baseline, sorted by SpatDistSpear — Part A: log-ratio metrics (ideal = 0).

| Method | LenDistort | CurvRatio | SegVarRatio | SpatSimRatio |
| --- | --- | --- | --- | --- |
| CNPE | 0.701 | 0.376 | -0.820 | 0.261 |
| LPMIP | 0.636 | 0.267 | -0.756 | 0.270 |
| CISOMAP | 1.393 | 0.283 | 0.491 | 0.685 |
| SPE | 0.518 | 0.122 | -1.463 | 0.210 |
| MDS | 0.890 | 0.105 | -1.067 | 0.464 |
| DM | 2.721 | 0.354 | 2.322 | 1.144 |
| PHATE | 2.976 | 0.736 | 1.932 | 1.174 |
| DVE | 4.563 | 0.382 | 5.322 | 1.377 |
| LAPEIG | 4.462 | 0.338 | 5.147 | 1.372 |
| SPMDS | 3.102 | 0.318 | 2.176 | 1.207 |
| UMAP | 0.160 | 0.515 | -4.550 | 0.218 |
| TSNE | -0.035 | 0.346 | -4.395 | 0.008 |
| FOSMOD | 1.216 | 0.121 | -0.315 | 0.775 |
| PFA | 1.050 | -0.070 | -0.532 | 0.756 |
| NNP | -0.202 | -0.070 | -3.111 | 0.047 |
| LPP | 1.289 | -0.373 | -0.783 | 1.063 |

**Table 10:**
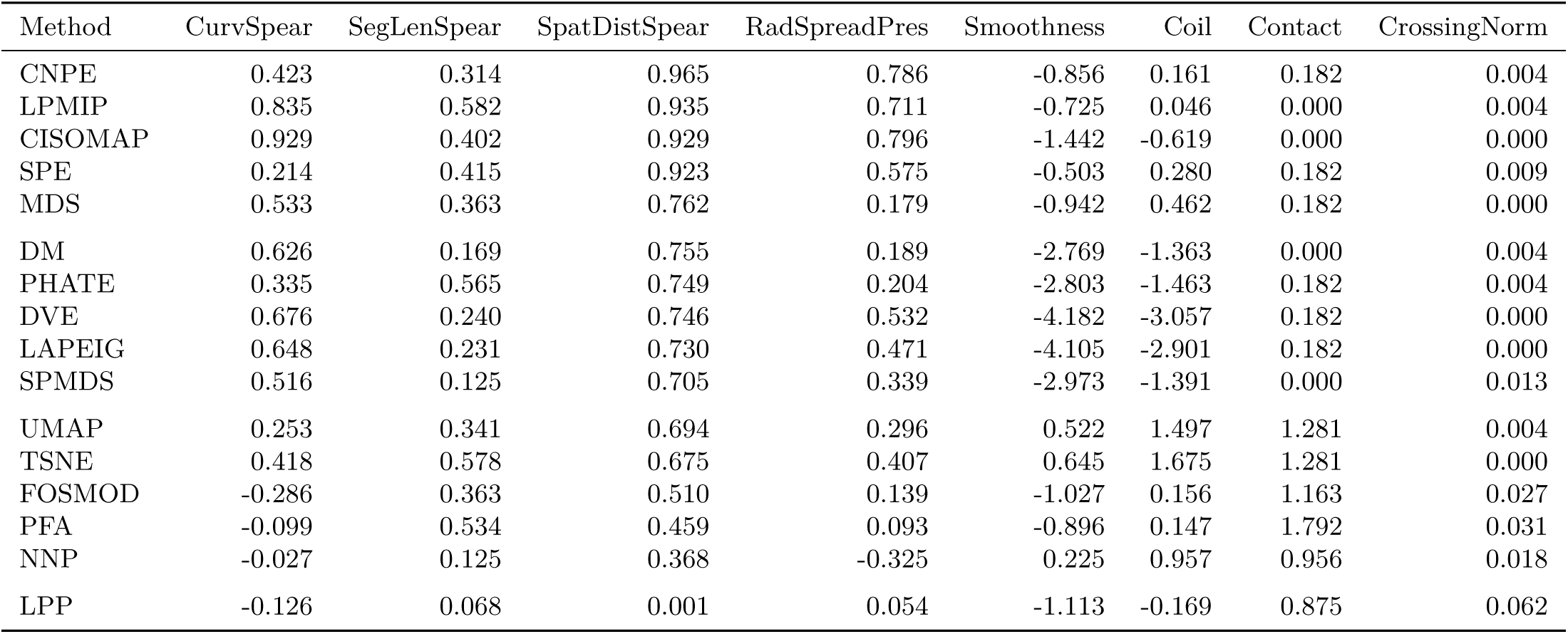
Cyclic-loop preservation metrics at the 15 nodes baseline — Part B: Spearman preservation metrics (ideal = 1); RadSpreadPres (ideal = 1); Smoothness and Contact (ideal = low); Coil (ideal = high); CrossingNorm (ideal = low).

| Method | CurvSpear | SegLenSpear | SpatDistSpear | RadSpreadPres | Smoothness | Coil | Contact | CrossingNorm |
| --- | --- | --- | --- | --- | --- | --- | --- | --- |
| CNPE | 0.423 | 0.314 | 0.965 | 0.786 | -0.856 | 0.161 | 0.182 | 0.004 |
| LPMIP | 0.835 | 0.582 | 0.935 | 0.711 | -0.725 | 0.046 | 0.000 | 0.004 |
| CISOMAP | 0.929 | 0.402 | 0.929 | 0.796 | -1.442 | -0.619 | 0.000 | 0.000 |
| SPE | 0.214 | 0.415 | 0.923 | 0.575 | -0.503 | 0.280 | 0.182 | 0.009 |
| MDS | 0.533 | 0.363 | 0.762 | 0.179 | -0.942 | 0.462 | 0.182 | 0.000 |
| DM | 0.626 | 0.169 | 0.755 | 0.189 | -2.769 | -1.363 | 0.000 | 0.004 |
| PHATE | 0.335 | 0.565 | 0.749 | 0.204 | -2.803 | -1.463 | 0.182 | 0.004 |
| DVE | 0.676 | 0.240 | 0.746 | 0.532 | -4.182 | -3.057 | 0.182 | 0.000 |
| LAPEIG | 0.648 | 0.231 | 0.730 | 0.471 | -4.105 | -2.901 | 0.182 | 0.000 |
| SPMDS | 0.516 | 0.125 | 0.705 | 0.339 | -2.973 | -1.391 | 0.000 | 0.013 |
| UMAP | 0.253 | 0.341 | 0.694 | 0.296 | 0.522 | 1.497 | 1.281 | 0.004 |
| TSNE | 0.418 | 0.578 | 0.675 | 0.407 | 0.645 | 1.675 | 1.281 | 0.000 |
| FOSMOD | -0.286 | 0.363 | 0.510 | 0.139 | -1.027 | 0.156 | 1.163 | 0.027 |
| PFA | -0.099 | 0.534 | 0.459 | 0.093 | -0.896 | 0.147 | 1.792 | 0.031 |
| NNP | -0.027 | 0.125 | 0.368 | -0.325 | 0.225 | 0.957 | 0.956 | 0.018 |
| LPP | -0.126 | 0.068 | 0.001 | 0.054 | -1.113 | -0.169 | 0.875 | 0.062 |

**Table 11:** Largest 20 changes between the 15 nodes and 10% paths (sorted by |Delta|).

| Method | metric | V_base | V_wide | Delta |
| --- | --- | --- | --- | --- |
| UMAP | SegVarRatio | -4.550 | -1.564 | 2.987 |
| SPMDS | SegVarRatio | 2.176 | 5.065 | 2.888 |
| PHATE | SegVarRatio | 1.932 | 4.471 | 2.539 |
| TSNE | SegVarRatio | -4.395 | -2.250 | 2.145 |
| LAPEIG | SegVarRatio | 5.147 | 7.288 | 2.140 |
| LPP | SegVarRatio | -0.783 | 1.307 | 2.090 |
| DVE | SegVarRatio | 5.322 | 7.412 | 2.089 |
| TSNE | Contact | 1.281 | 0.029 | -1.252 |
| UMAP | Contact | 1.281 | 0.035 | -1.246 |
| MDS | SegVarRatio | -1.067 | 0.154 | 1.221 |
| DM | SegVarRatio | 2.322 | 3.482 | 1.159 |
| PFA | SegVarRatio | -0.532 | 0.623 | 1.156 |
| NNP | SegVarRatio | -3.111 | -1.957 | 1.154 |
| SPE | SegVarRatio | -1.463 | -0.334 | 1.130 |
| UMAP | Coil | 1.497 | 0.474 | -1.023 |
| PFA | Contact | 1.792 | 0.777 | -1.014 |
| LPMIP | SegVarRatio | -0.756 | 0.211 | 0.968 |
| CNPE | SegVarRatio | -0.820 | 0.145 | 0.965 |
| FOSMOD | SegVarRatio | -0.315 | 0.637 | 0.952 |
| LAPEIG | Coil | -2.901 | -3.798 | -0.897 |

**Table 12:** Mean and maximum absolute change per metric between the 15 nodes and 10% paths, across all methods.

| Metric | MeanAbsDelta | MaxAbsDelta |
| --- | --- | --- |
| SegVarRatio | 1.648 | 2.987 |
| Coil | 0.448 | 1.023 |
| Contact | 0.380 | 1.252 |
| RadSpreadPres | 0.324 | 0.633 |
| Smoothness | 0.276 | 0.757 |
| CurvRatio | 0.275 | 0.696 |
| CurvSpear | 0.258 | 0.609 |
| LenDistort | 0.249 | 0.736 |
| SegLenSpear | 0.174 | 0.480 |
| SpatDistSpear | 0.123 | 0.349 |
| SpatSimRatio | 0.090 | 0.224 |
| CrossingNorm | 0.013 | 0.037 |

**Table 13:** Per-method SD across density levels — Part A: log-ratio metrics. Methods sorted by overall sensitivity (most stable first).

| Method | LenDistort | CurvRatio | SegVarRatio | SpatSimRatio |
| --- | --- | --- | --- | --- |
| SPE | 0.060 | 0.051 | 0.350 | 0.007 |
| CNPE | 0.038 | 0.107 | 0.321 | 0.022 |
| LPMIP | 0.066 | 0.073 | 0.320 | 0.019 |
| CISOMAP | 0.046 | 0.075 | 0.270 | 0.016 |
| MDS | 0.054 | 0.048 | 0.405 | 0.009 |
| DM | 0.040 | 0.104 | 0.393 | 0.024 |
| FOSMOD | 0.065 | 0.042 | 0.322 | 0.026 |
| NNP | 0.036 | 0.056 | 0.363 | 0.068 |
| PFA | 0.033 | 0.040 | 0.356 | 0.019 |
| LAPEIG | 0.165 | 0.089 | 0.691 | 0.039 |
| DVE | 0.148 | 0.102 | 0.664 | 0.040 |
| LPP | 0.090 | 0.105 | 0.673 | 0.039 |
| PHATE | 0.187 | 0.214 | 0.777 | 0.033 |
| SPMDS | 0.221 | 0.102 | 0.896 | 0.040 |
| TSNE | 0.154 | 0.120 | 0.671 | 0.074 |
| UMAP | 0.220 | 0.142 | 0.892 | 0.046 |

**Table 14:** Per-method SD across density levels — Part B: Spearman preservation and shape metrics.

| Method | CurvSpear | SegLenSpear | SpatDistSpear | RadSpreadPres | Smoothness | Coil | Contact |
| --- | --- | --- | --- | --- | --- | --- | --- |
| SPE | 0.082 | 0.087 | 0.009 | 0.093 | 0.058 | 0.056 | 0.044 |
| CNPE | 0.065 | 0.113 | 0.003 | 0.045 | 0.051 | 0.103 | 0.048 |
| LPMIP | 0.137 | 0.059 | 0.006 | 0.045 | 0.034 | 0.066 | 0.148 |
| CISOMAP | 0.166 | 0.074 | 0.019 | 0.045 | 0.061 | 0.102 | 0.102 |
| MDS | 0.089 | 0.079 | 0.047 | 0.198 | 0.089 | 0.072 | 0.123 |
| DM | 0.104 | 0.163 | 0.052 | 0.185 | 0.073 | 0.075 | 0.116 |
| FOSMOD | 0.188 | 0.039 | 0.084 | 0.188 | 0.041 | 0.146 | 0.249 |
| NNP | 0.116 | 0.092 | 0.030 | 0.149 | 0.027 | 0.138 | 0.131 |
| PFA | 0.146 | 0.090 | 0.100 | 0.197 | 0.041 | 0.114 | 0.316 |
| LAPEIG | 0.076 | 0.074 | 0.034 | 0.073 | 0.198 | 0.281 | 0.087 |
| DVE | 0.083 | 0.085 | 0.058 | 0.066 | 0.204 | 0.239 | 0.079 |
| LPP | 0.110 | 0.104 | 0.029 | 0.084 | 0.117 | 0.258 | 0.274 |
| PHATE | 0.071 | 0.101 | 0.047 | 0.187 | 0.160 | 0.182 | 0.058 |
| SPMDS | 0.054 | 0.168 | 0.059 | 0.136 | 0.173 | 0.212 | 0.124 |
| TSNE | 0.141 | 0.112 | 0.013 | 0.132 | 0.193 | 0.242 | 0.361 |
| UMAP | 0.117 | 0.093 | 0.038 | 0.145 | 0.243 | 0.343 | 0.403 |

**Table 15:** Overall sensitivity of each method to loop density, ranked from most stable (rank 1) to least stable. Score = mean rescaled SD across all 12 cyclic metrics.

| Rank | Method | OverallSensitivity |
| --- | --- | --- |
| 1 | SPE | 0.124 |
| 2 | CNPE | 0.148 |
| 3 | LPMIP | 0.151 |
| 4 | CISOMAP | 0.200 |
| 5 | MDS | 0.293 |
| 6 | DM | 0.351 |
| 7 | FOSMOD | 0.371 |
| 8 | NNP | 0.377 |
| 9 | PFA | 0.382 |
| 10 | LAPEIG | 0.406 |
| 11 | DVE | 0.419 |
| 12 | LPP | 0.472 |
| 13 | PHATE | 0.516 |
| 14 | SPMDS | 0.538 |
| 15 | TSNE | 0.591 |
| 16 | UMAP | 0.682 |

#### 3.2.3 Non-parametric rank stability across thresholds

As for the linear path, we summarise each method’s performance across all metrics and all density levels with a composite dense rank, and display the distribution as a beanplot. Direction conventions are adapted to the cyclic metric set: the four log-ratio metrics have ideal 0, the four Spearman preservation metrics have ideal 1, `Smoothness`, `Contact`, and `CrossingNorm` are best when low, and `Coil` is best when high.

In the cyclic loop, the methods concentrated at the best (lowest) composite ranks are CNPE, LPMIP, SPE, MDS, while those concentrated at the worst ranks are DVE, LPP, LAPEIG, SPMDS. This ordering largely echoes the linear case at both ends of the spectrum: CNPE, LPMIP, SPE, and MDS are robust top performers on both the linear path and the closed loop, and LAPEIG, DVE, and SPMDS are consistently poor performers on both. UMAP and TSNE are the exception — both rank in the top six on the linear PC1 path but drop out of the top tier on the closed loop, falling to the middle of the sixteen-method panel rather than to the bottom, which suggests that these neighbour-based embeddings preserve a linear axis more reliably than a closed cycle without becoming actively poor performers on it. Despite this partial reordering of *which* methods do best, the within-dataset conclusion is the same as for the linear path: each method’s composite rank is stable across all density levels, so the loop’s benchmark ordering is not driven by the path-density threshold.

#### 3.2.4 Projected loop by density level

### 3.3 15 nodes (n = 15)

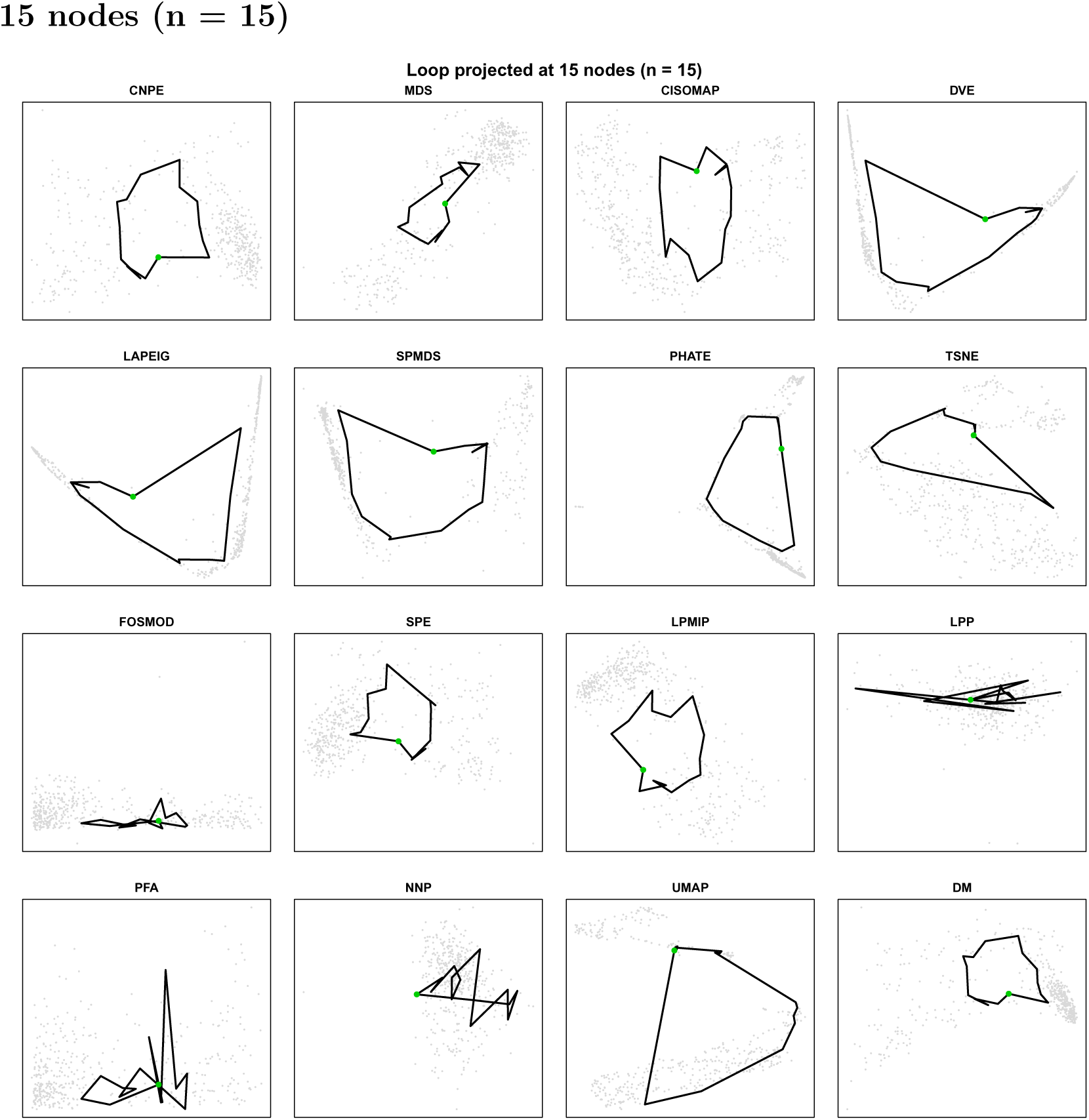

### 3.4 1% (n = 38)

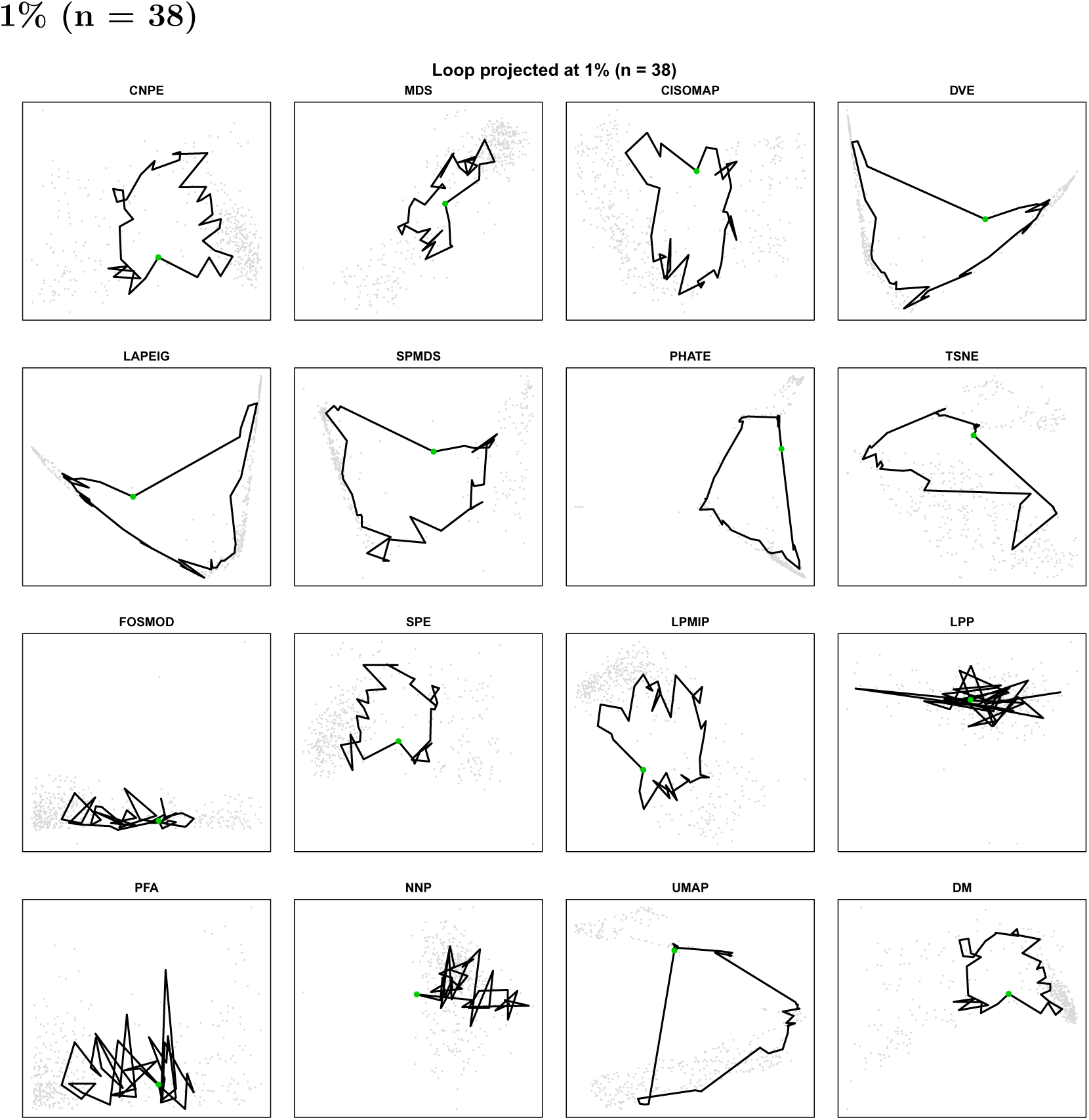

### 3.5 2% (n = 62)

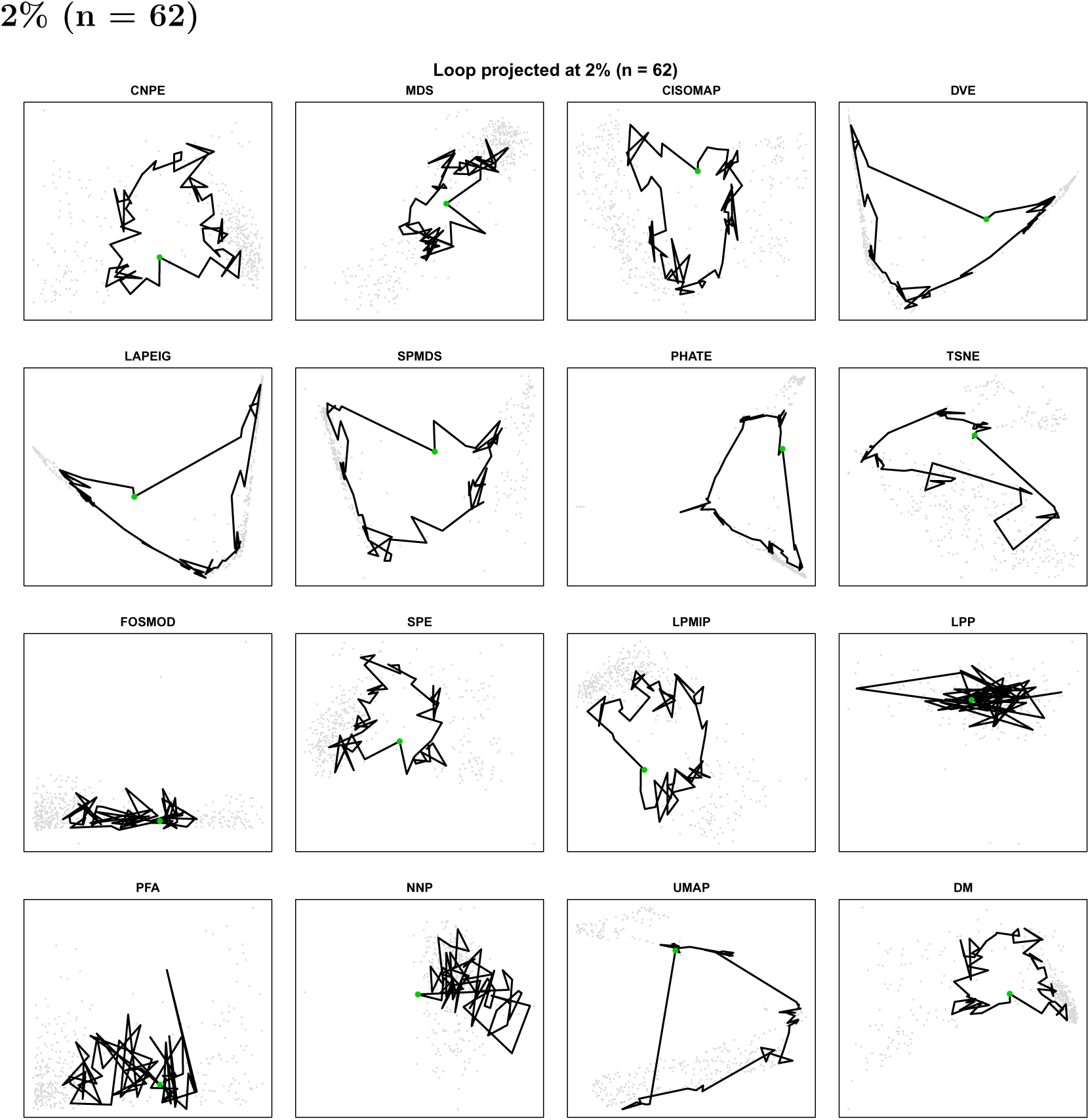

### 3.6 3% (n = 87)

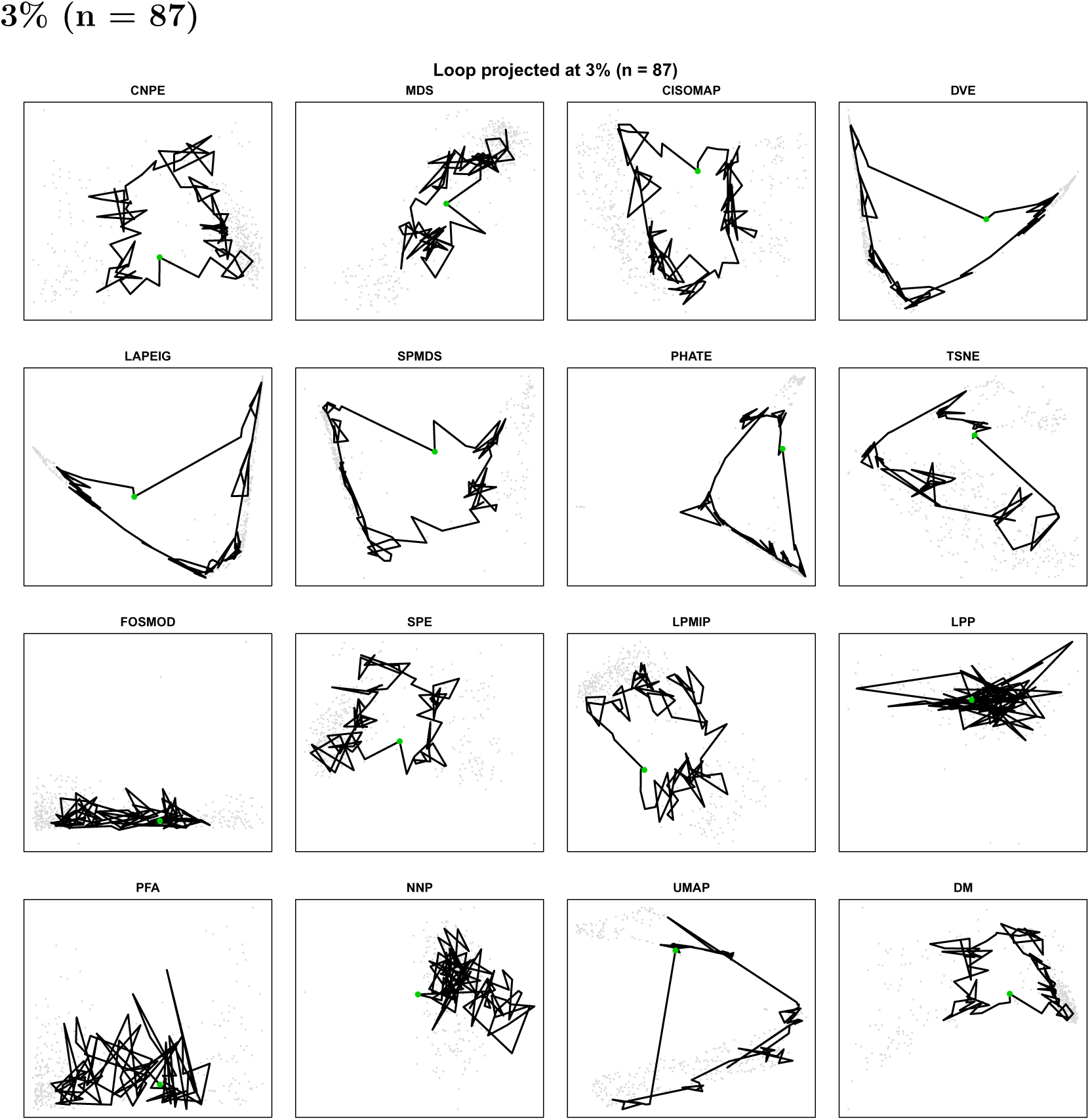

### 3.7 4% (n = 102)

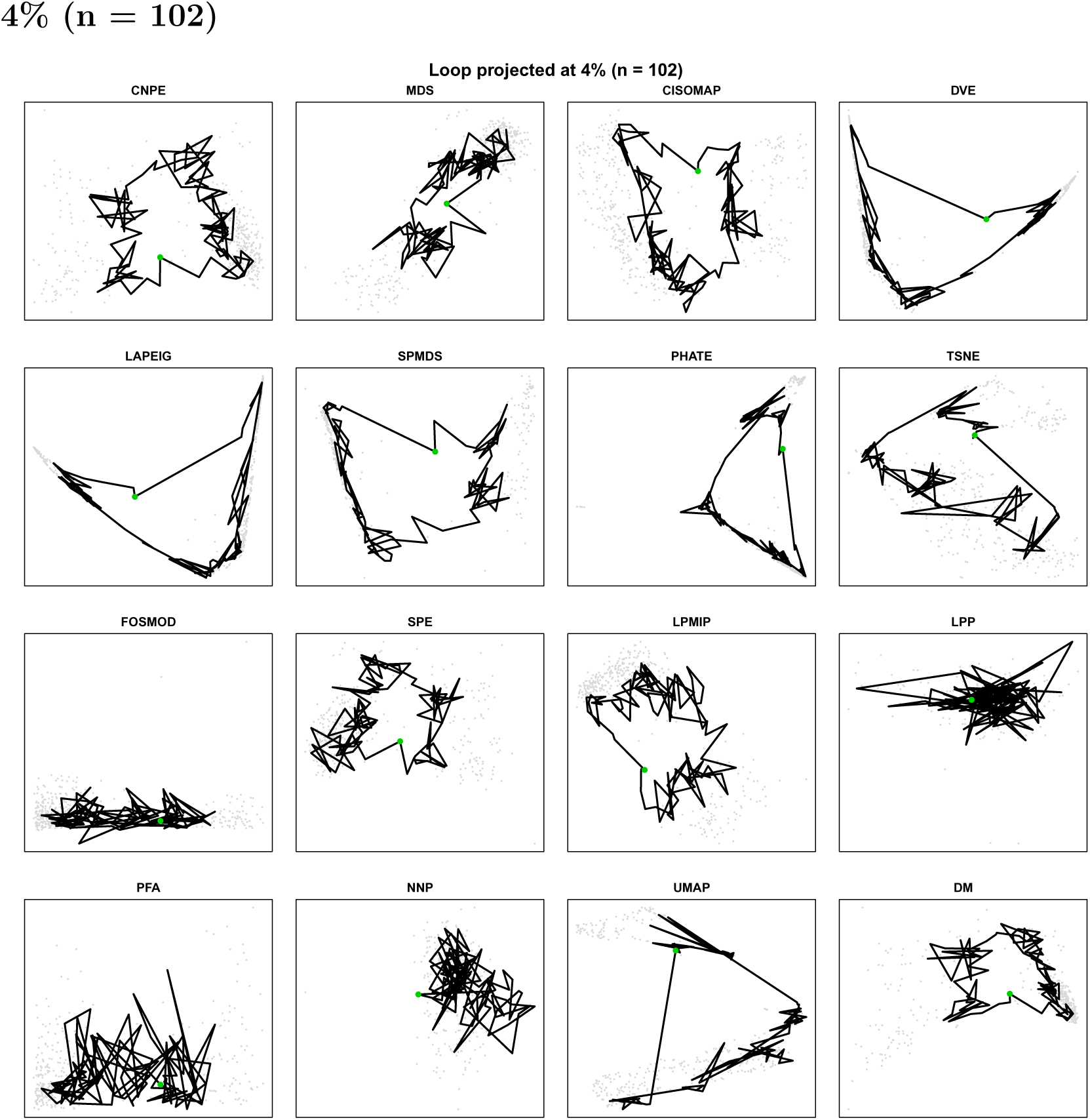

### 3.8 5% (n = 122)

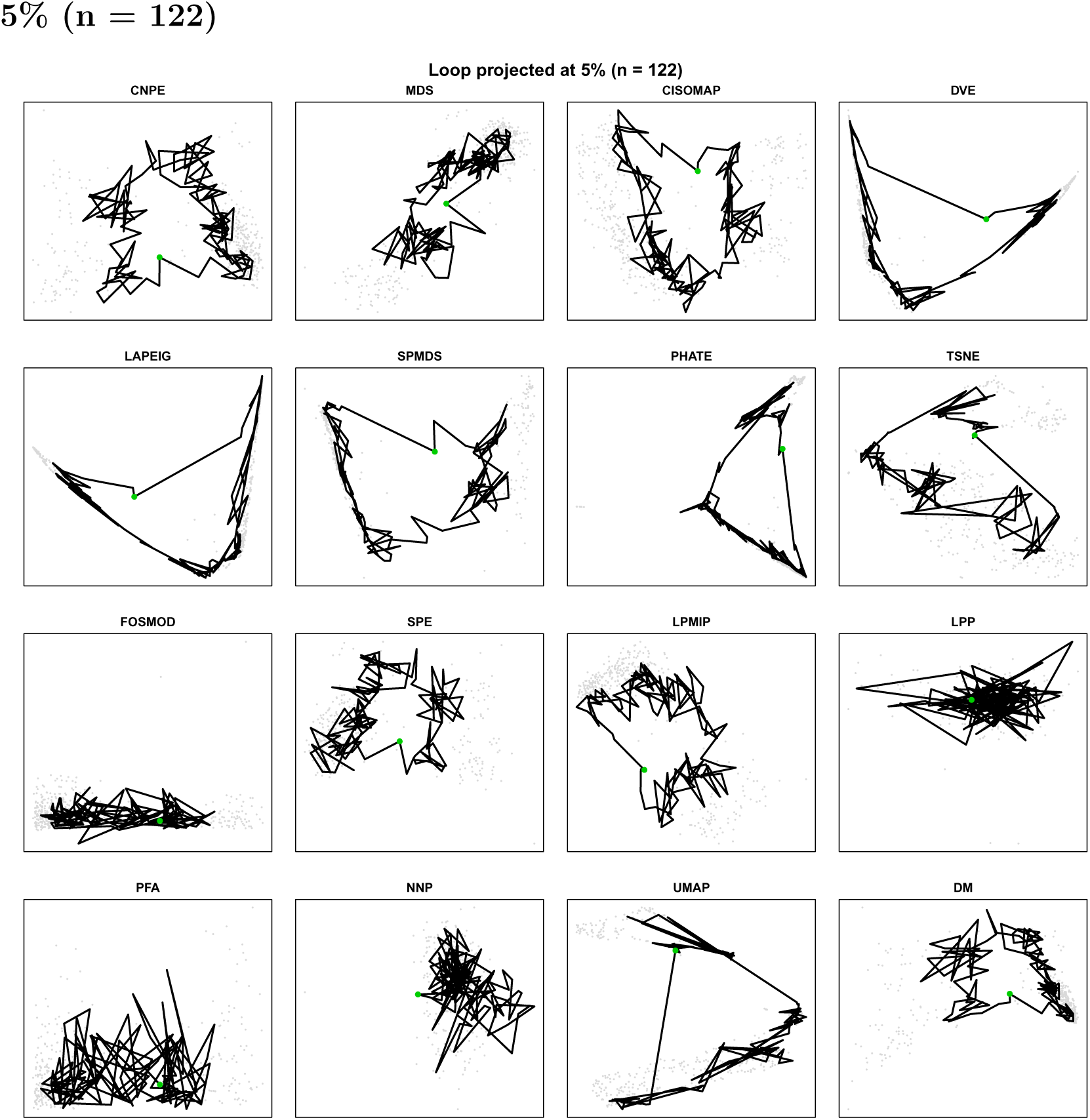

### 3.9 6% (n = 141)

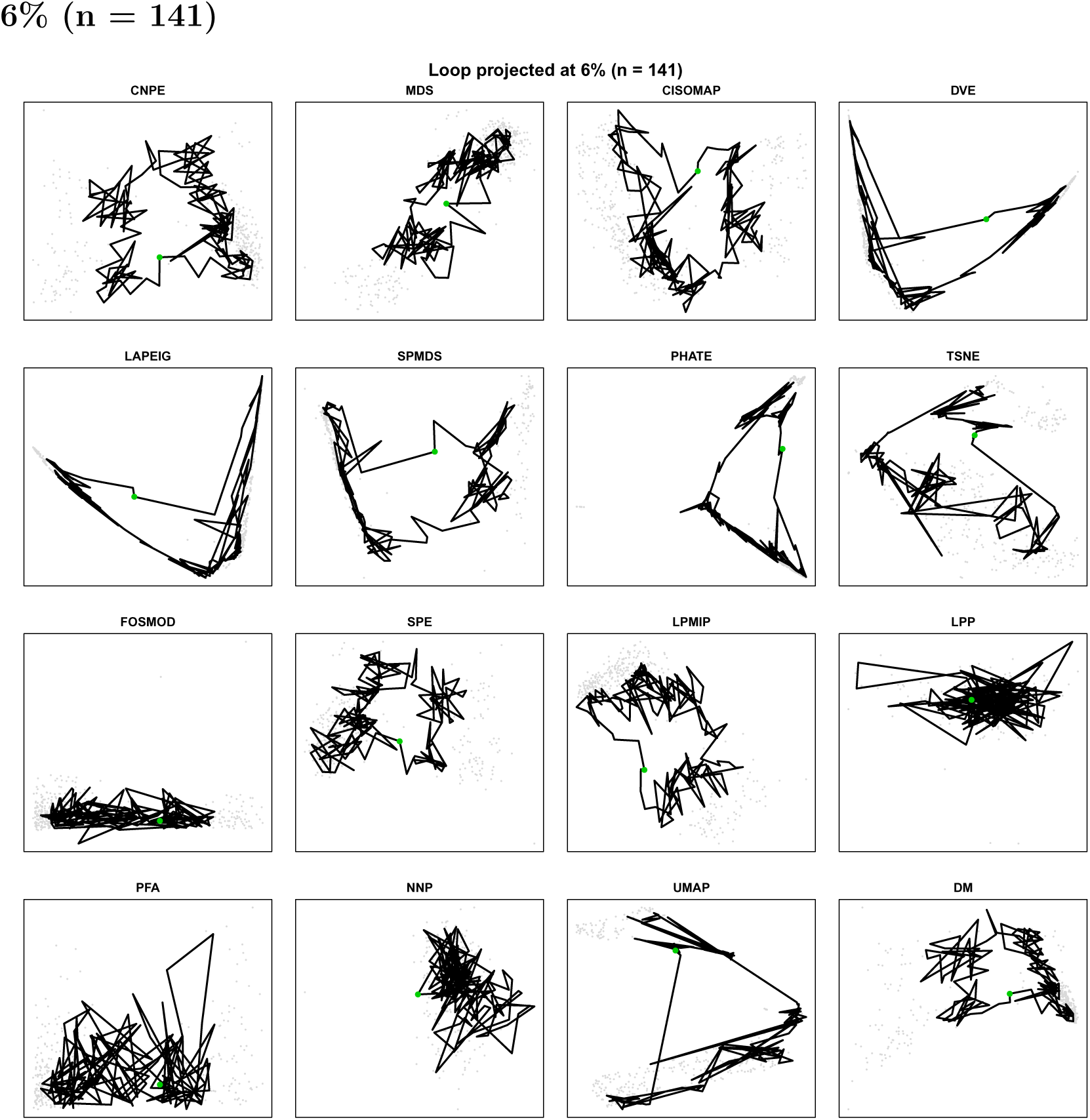

### 3.10 7% (n = 157)

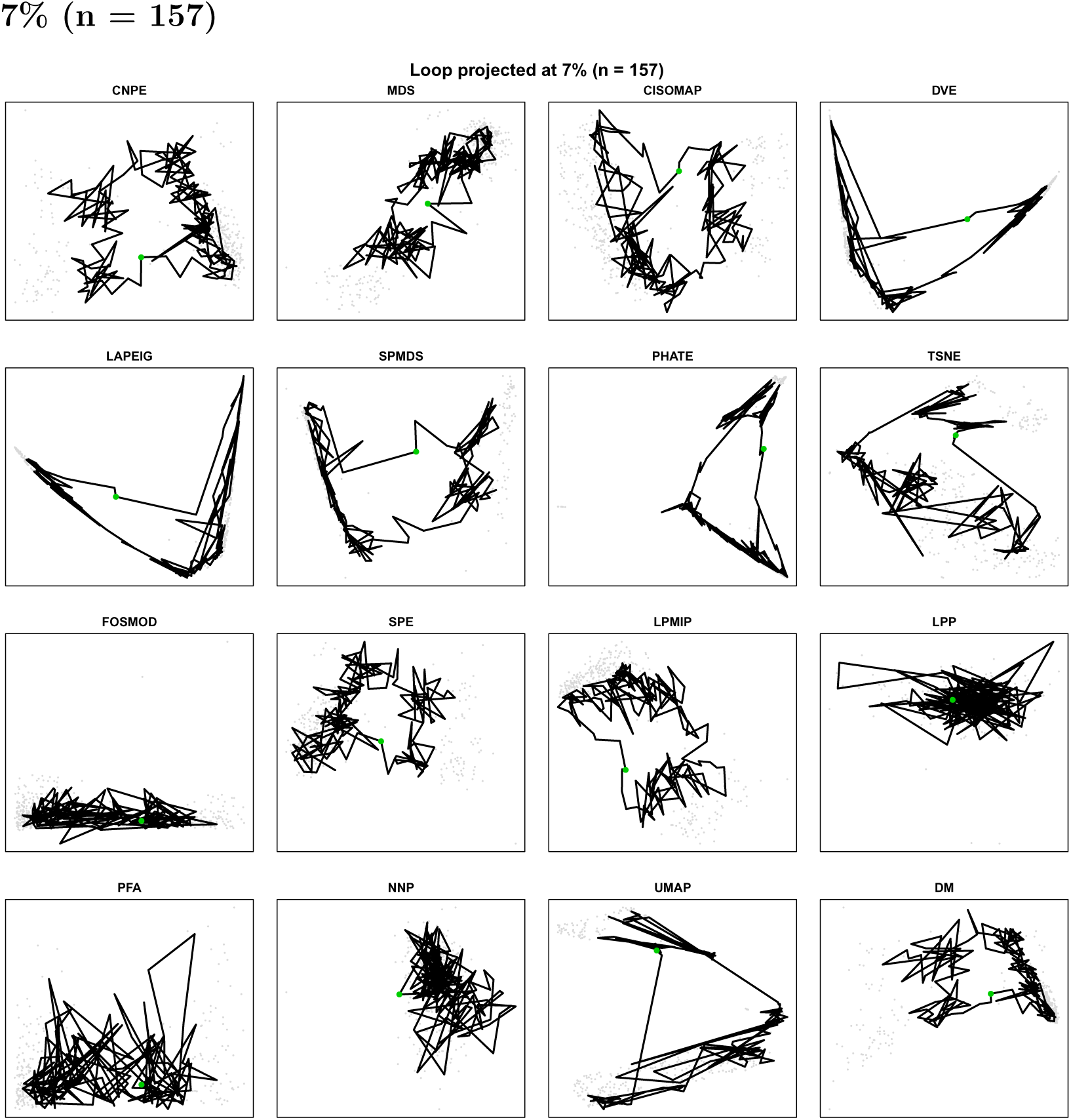

### 3.11 8% (n = 170)

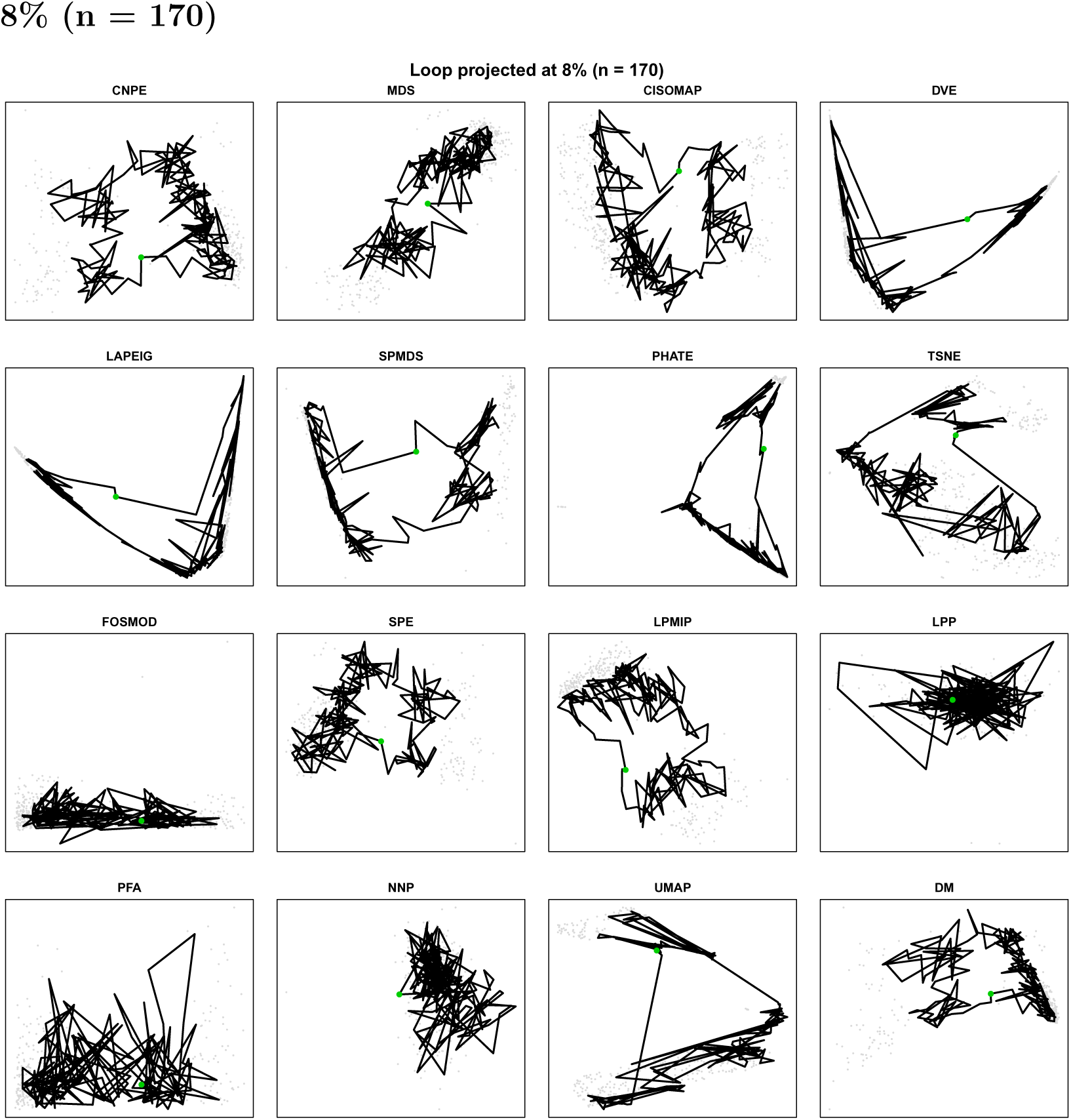

### 3.12 9% (n = 183)

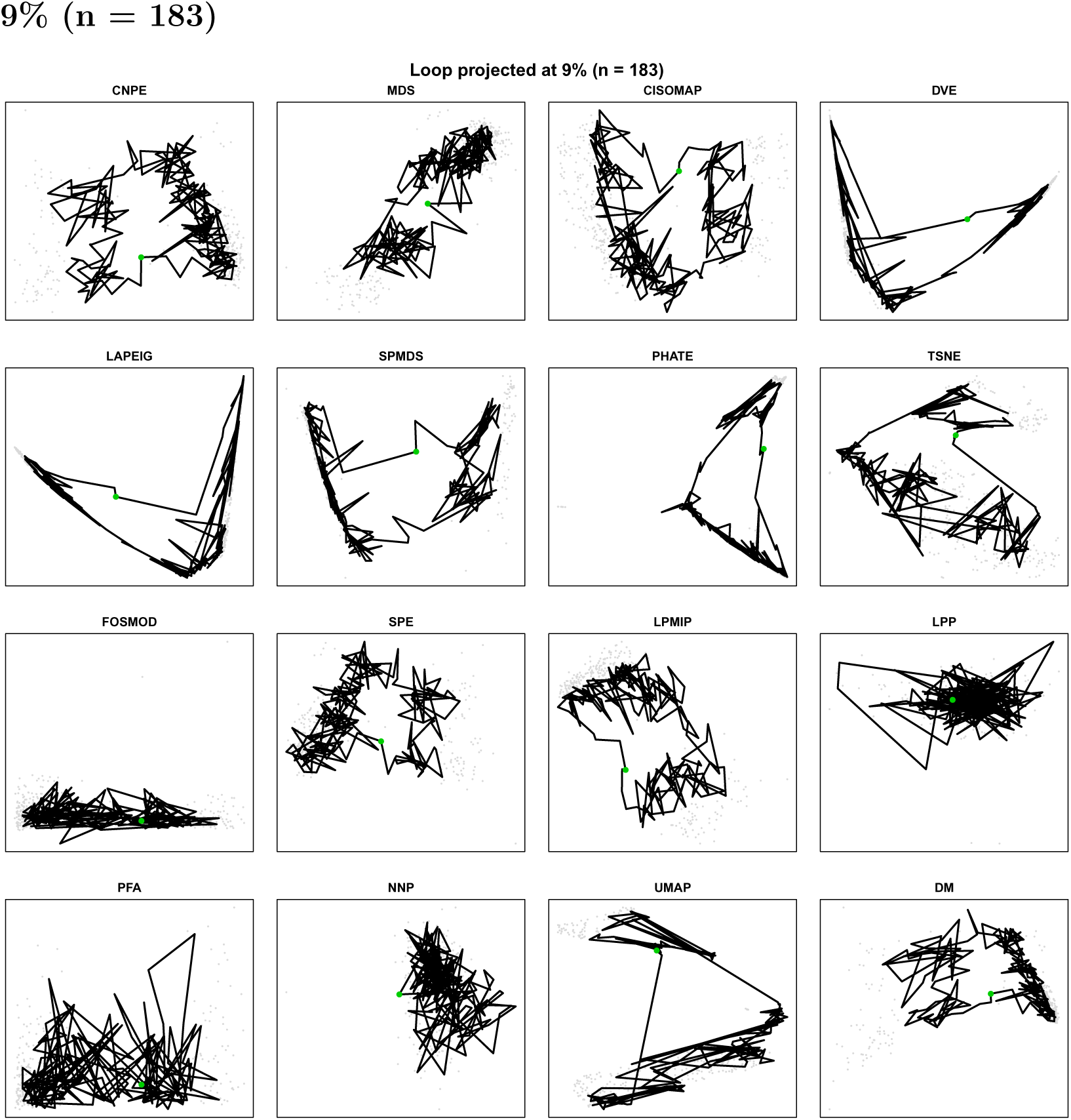

### 3.13 10% (n = 195)

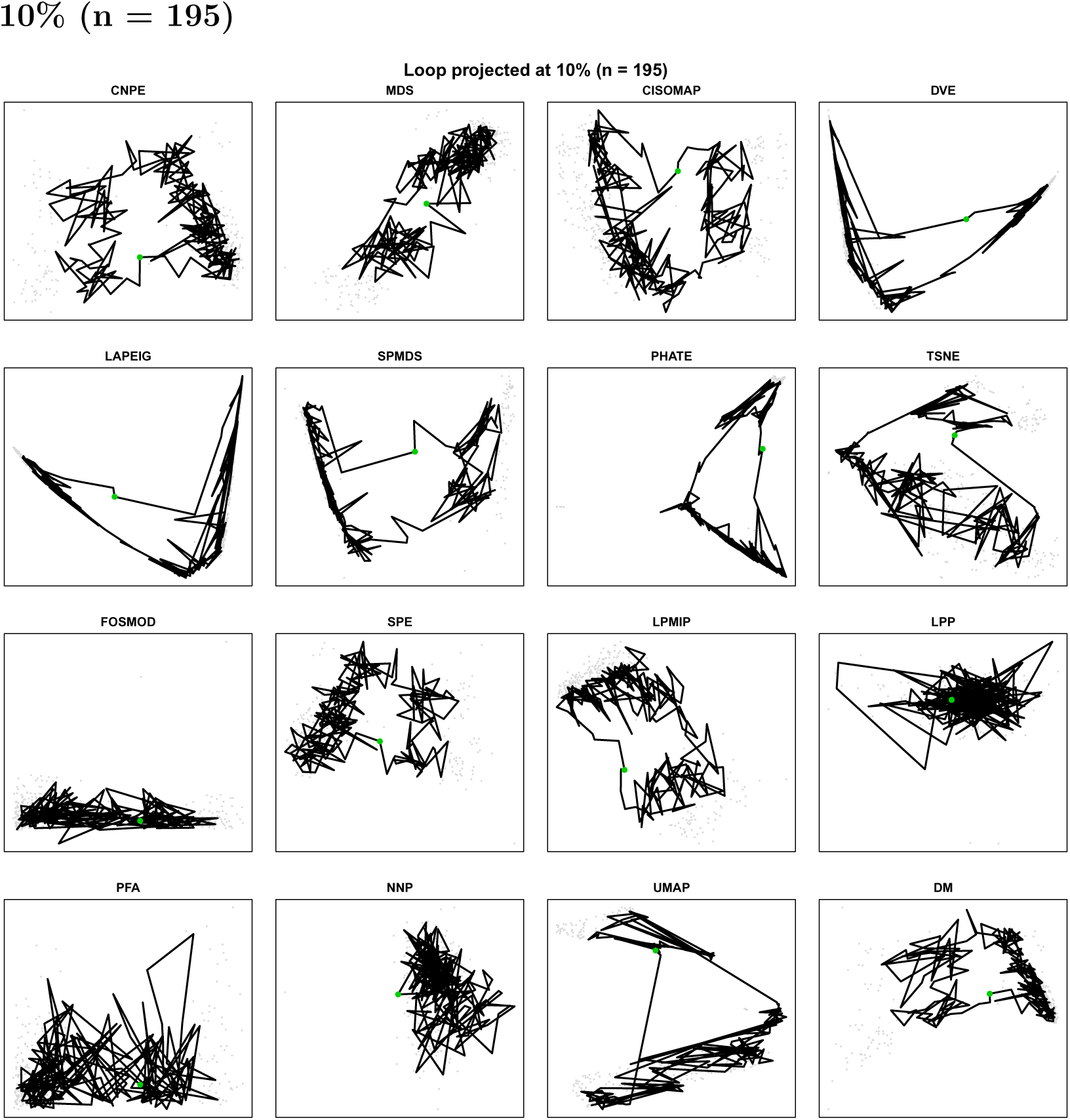

Going from the 15 nodes baseline to the widest 10% band (195 ordered points), the most density-**stable** method is **SPE** and the most density-**sensitive** is **UMAP**. The single largest metric change is **SegVarRatio** for **UMAP** (-4.55 → -1.564, Δ 2.987), and across methods the metric that moves most on average is **SegVarRatio**.

As in the linear case, the absolute metric values respond to loop density while the relative method ordering is preserved: widening the per-segment band from the 15-node baseline to the 10% level pulls in cells farther from the loop polyline, shifting the log-ratio and Spearman metrics in parallel without reordering the methods. The shape metrics (`Smoothness`, `Coil`, `Contact`) from the Preservation package confirm that methods which embed the loop as a smooth, well-separated cycle remain the same across density levels. The three metric families — log-ratios, Spearman preservations, and Preservation-package shape metrics — broadly agree on the method ordering, mirroring the consensus seen for the linear path. This consistency across a topologically distinct reference supports the conclusion that path-density threshold choice does not drive the benchmark rankings.

## 4 Discussion

This analysis demonstrates that path-preservation benchmarks are robust to the choice of density threshold across the 1–10% range examined, for both a linear PC1-proximity reference path and a closed cell-cycle loop. Absolute metric values — particularly `SegVarRatio`, `Smoothness`, and the Spearman correlation metrics — shift as the threshold grows, but these shifts are largely parallel across methods and reflect properties of the reference path itself rather than changes in embedding quality. Specifically, as more peripheral cells are included, the reference path broadens and becomes less strictly aligned with its underlying geometry, introducing mild distortion that affects all methods simultaneously. The key finding, observed identically in the linear and cyclic analyses, is that this systematic drift does not disrupt the relative ordering: the composite dense rank of each method is stable across all density levels, and no method changes performance tier — even though the absolute per-method sensitivity ranking itself differs between the two reference geometries.

The threshold-sensitivity results have a practical implication for the design of DR benchmarks. The 10% threshold used as the operating point in the parent framework is not a critical parameter choice — values anywhere from 2% to 10% would yield the same conclusions about which methods preserve the path well. We recommend against using 1% in practice, not because it changes the rankings, but because 31 cells is a sufficiently small sample that rank-correlation metrics become noisy; this is visible in the CurvSpear and SegLenSpear panels of Figure∼5, which show wider inter-method spread at 1% than at 3–10%, while the same panels are stable from 3% onward. Structural complexity metrics can also be affected by discretisation artefacts at very low cell counts. A threshold of 5–10% (155–310 cells) provides a stable operating region.

A secondary finding concerns the relationship between threshold-sensitivity and performance quality. Among the most threshold-stable methods are CNPE (0.27) and, notably, FOSMOD (0.30) and UMAP (0.34), some of which are otherwise modest path-preservers. Conversely, CISOMAP (0.52) is a strong overall performer but among the most threshold-sensitive, suggesting that its metric values are more strongly influenced by path composition than other high-quality methods. This decoupling between quality and stability means that threshold-stability alone is not a useful criterion for method selection; the absolute performance level at any fixed threshold remains the appropriate primary criterion.

Several limitations apply to this analysis. The study uses a single cluster from a single CyTOF dataset; whether the conclusions generalise to scRNA-seq data, to non-linear trajectories, or to clusters where PC1 is not the dominant axis of variation remains to be established. The reference path is defined by proximity to a linear axis and therefore inherently favours methods that preserve global linear structure; methods optimised for local neighbourhood preservation (UMAP, t-SNE) may be disadvantaged by this construction relative to their performance on non-linear manifolds. We addressed the linearity concern directly by repeating the analysis on a closed cell-cycle loop (Section 3.2); the rank-stability conclusion held there as well, indicating that the robustness to path density is not specific to linear trajectories. Across both the linear CD4^+^ T-cell path and the cyclic B-cell loop, absolute metric values responded to path density while the method rankings did not, which is the central, dataset-independent result of this work. Extension to scRNA-seq data and to branching trajectories remains a natural follow-up.

## 5 Data and Code Availability

The operational validation configurations, parameter data structures, and non-parametric ranking functions utilized to assess threshold stability are open-source and accessible at https://github.com/pbombina/path-preservation.

## Notes

### Competing Interest Statement

The authors have declared no competing interest.

